# An optogenetics-compatible red fluorescent calcium indicator with negligible blue light photoactivation

**DOI:** 10.64898/2026.02.28.708321

**Authors:** Xueling Zhang, Brittany R. Addison, Emine Zeynep Ulutas, Claire M. Deng, Samara Doshi, Sydney Nabhan, Alan J. Emanuel, Jeffrey E. Markowitz, Dorothy Koveal

## Abstract

Red genetically encoded calcium indicators (GECIs) are important tools for live cell and *in vivo* imaging. However, their application in optogenetic experiments has been limited by their complex photophysics, which can yield blue-light-induced artifacts. These photophysical drawbacks arise from the fluorescent protein (FP) used to construct the GECI. To address these limitations, we engineered novel red GECIs based on photostable red FPs, including mScarlet variants. After testing multiple design topologies and screening for calcium responses, we identified a lead variant, named ScaRCaMP-1.0. ScaRCaMP-1.0 exhibits moderate Ca^2+^ responses (ΔF/F_0_ = -13%) relative to other red GECIs, a tradeoff that appears to have enabled remarkable blue-light photostability at power densities exceeding 200 mW/mm^2^. We validated the performance of ScaRCaMP-1.0 in an optogenetic regime, and *in vivo* via fiber photometry. Finally, guided by structural predictions, we investigated the mechanism underlying ScaRCaMP responses. A pair of lysine residues on the surface of the FP appear to be important for controlling Ca^2+^ responses, and mutation of one residue (K132Y) notably increased the response size (ΔF/F_0_ = -22%) without compromising blue-light photostability. We call the improved variant ScaRCaMP-2.0. Taken together, these results establish ScaRCaMP as an optogenetics-compatible red GECI and demonstrate the potential of mScarlet-based fluorophores as a basis for generating photostable red biosensors.

## Introduction

Calcium imaging is a foundational approach for measuring neuronal activity in the brain. Genetically encoded calcium indicators (GECIs) detect intracellular changes in free calcium ions (Ca^2+^) that accompany cell signaling events or an action potential. GECIs harness Ca^2+^-induced conformational changes between a Ca^2+^ binding domain, calmodulin (CaM), and its cognate peptide to induce fluorescence changes in a fused fluorescent protein.^1–3^ GECIs can be readily deployed *in vivo* and targeted to specific brain regions, cell types, and subcellular locations, making them instrumental tools for dissecting neural pathways and circuits in live animals.^2–6^ GECIs are also available in a full palette of colors,^3,7,8^ making them spectrally compatible with other genetically encoded fluorescent biosensors and optical tools.

Equally important to reading neural activity is the ability to write neural activity. Optogenetic tools, including the growing lineage of channelrhodopsins,^9–13^ enable direct and precise neural activation or inhibition with pulses of blue light. Optogenetic tools have been combined with GECIs to map neural circuits and measure circuit dynamics *in vivo*.^14–17^ However, there remain limitations as to how these two classes of tools can be combined. First, optogenetic tools cannot easily be combined with green fluorescent GECIs due to spectral crosstalk.^15^ Green fluorescent GECIs strongly absorb the blue light used to activate optogenetic tools.^3,18,19^ Additionally, many red-shifted optogenetic tools have significant photocurrents at wavelengths used to excite green fluorescent GECIs.^20,21^ Furthermore, in green GECI/red opsin all-optical experiments, even if imaging light is insufficient to drive suprathreshold photocurrents, it can desensitize opsins and contaminate experimental results.^22^

Second, red fluorescent GECIs exhibit reversible photoswitching which can introduce spectral artifacts.^3^ During photoswitching, high-energy wavelengths in the 400–490 nm range induce fluorescence changes via a process that is distinct from fluorescence excitation. Photoswitching artifacts have limited the co-deployment of red GECIs with optogenetic tools in the same cells or brain region, except in cases where the activating light can be used at low power levels.^14,23,24^ Photoswitching artifacts can exhibit kinetics and amplitudes that mimic legitimate biosensor changes^25,26^ and their magnitude scales with illumination power, making them a general concern for red fluorescent biosensors.^25–34^

Photoswitching behavior arises from the underlying fluorescent protein (FP). Of the FPs used to construct GECIs and other biosensors, this property has been most evident in mApple;^35,36^ although photoswitching and other photochromic behaviors have also been observed in mRuby fluorophores.^36,37^ Photoswitching has been thoroughly characterized in mApple-based GECIs, such as jRGECO, as well as mApple-based genetically encoded fluorescent biosensors that detect other ligands.^25–34^ In addition, while mRuby-based jRCaMP1a and jRCaMP1b were not originally reported to photoswitch,^38^ we observe here that exposure to higher power blue light does induce moderate photoswitching consistent with that previously observed in the underlying mRuby fluorophore.^36,37^

We therefore set out to develop a red GECI with negligible photoswitching that also retains the brightness and strong *in vivo* expression profiles of current gold standard GECIs. Because photoswitching behavior arises from the fluorophore, we selected red fluorescent proteins that do not photoswitch. The mScarlet lineage of fluorescent proteins (including mScarlet, mScarlet3, mScarlet-I, and mScarlet-I3) is bright, photostable, and expresses well in mammalian cells.^36,39,40^ An improved version of mCherry, called mCherry-XL, also has a high quantum yield and strong expression in mammalian cells.^41^

Here we explored the adaptation of mCherry-XL and mScarlet fluorescent proteins for use in red GECIs. We present a new blue-light-insensitive red GECI based on mScarlet-I3, called ScaRCaMP-1.0, that has high baseline brightness and an inverse-response to Ca^2+^ (ΔF/F_0_ = -13%). We characterized its photochromic behavior in relation to leading red GECIs (jRGECO1a, jRCaMP1a and jRCaMP1b) and the newly-developed PinkyCaMP,^42^ and found that ScaRCaMP-1.0 is highly resistant to strong pulses of blue light (>200 mW/mm^2^). We further demonstrate the compatibility of ScaRCaMP-1.0 with optogenetic regimes, and we test its performance *in vivo* using fiber photometry. Overall, while its response to Ca^2+^ is moderate relative to other red GECIs, we find that the tradeoff for blue-light photostability may make ScaRCaMP-1.0 a valuable tool in experimental regimes where blue light power cannot easily be reduced. This represents an important step toward the creation of optogenetic-compatible biosensors. Finally, guided by AlphaFold3^43^ structure predictions, we explore the mechanism underlying ScaRCaMP’s fluorescence changes. We identify two surface-exposed lysine residues within the mScarlet-I3 fluorophore that influence Ca^2+^ responses. This led us to identify an improved variant (K132Y) called ScaRCaMP-2.0, which has a ΔF/F_0_ = -22% and maintains the blue-light photostability of its predecessor. Collectively, this work presents ScaRCaMP-1.0 and ScaRCaMP-2.0 as new optogenetics-compatible red GECIs and establishes that mScarlet-based fluorophores can be used to develop photostable red GECIs.

## Results

### Incorporation of photostable red fluorescent proteins into classic GECI designs

Existing red GECIs such as jRGECO display photoswitching behavior whereby blue light reversibly converts the GECI to a different fluorescent state following a process that is distinct from photon absorption and excitation.^3,35,44^ The result is blue-light-induced fluorescence artifacts (**Fig. 1a**). Importantly, blue-light-induced photoswitching is understood to arise from the fluorescent protein used to construct the GECI, most notably mApple, but also mRuby-based fluorophores.^35–37^ Newer red fluorescent proteins (RFPs), such as mCherry-XL and the mScarlet lineage of RFPs, are resistant to blue light.^36,39–41^ We therefore explored their use in classical GECI designs.

**Figure 1.**
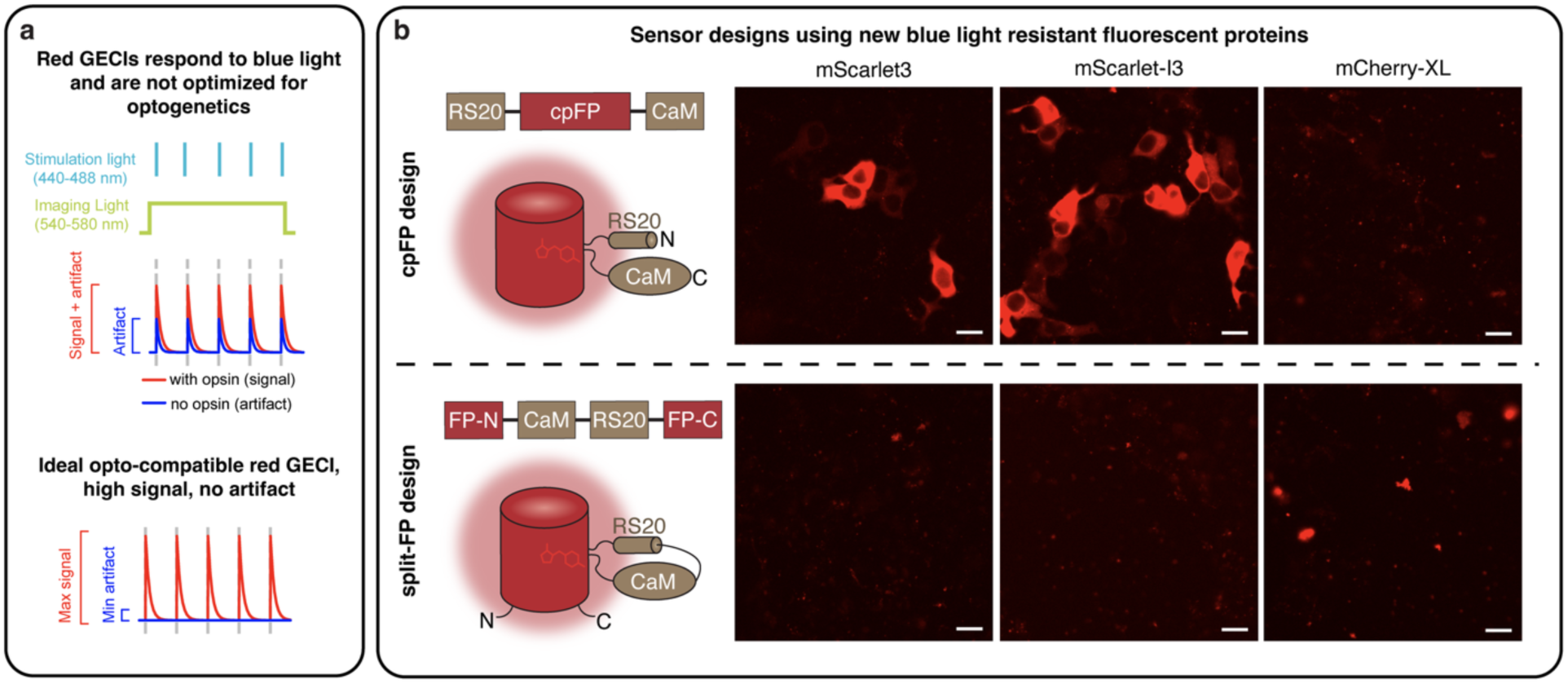
Making red GECIs that are compatible with optogenetic tools. (**a**) Schematic illustrating red GECI responses in optogenetic regimes. Existing red GECIs demonstrate blue light responses even in the absence of opsins. In an ideal scenario, the red GECI would show a response only in the presence of an opsin and would not respond to blue light alone. (**b**) We tested blue-light-resistant RFPs in two classic GECI designs: a circularly permuted fluorescent protein (cpFP)-based GECI and a split-FP-based GECI. Constructs were evaluated for their ability to be stably expressed in mammalian cells. HEK293T cells expressing initial GECIs designed using the fluorescent proteins indicated at the top, following the design schematic listed on the left. Scale bar represents 20 μm. RFP: red fluorescent protein, CaM: calmodulin, RS20: a calmodulin binding peptide.

Single-color GECIs comprise a split- or circularly permuted fluorescent protein (split-FP or cpFP) attached to the calcium-binding domain, calmodulin (CaM), and a Ca^2+^/CaM-binding peptide, such as the RS20 peptide.^2–4,7,45–53^ The archetypal cpFP GECIs, GCaMP and RCaMP, have been extensively optimized over multiple decades to achieve signals that are bright and fast, with exceptionally high signal-to-noise ratios *in vivo*.^2,4,38,49^ Split-FP versions^54–62^ such as NEMOf have also been made that circumvent the need for circular permutation of the fluorescent protein. Both design approaches have yielded bright and sensitive GECIs. We therefore created mScarlet3-, mScarlet-I3- and mCherry-XL-based designs following each topology (**Fig. 1b**). Circular permutation sites and split-FP sites were selected based on sequence homology with existing red GECIs.^3,23,38,63^ Linker sequences from jRCaMP1b^38^ were used in our initial cpFP designs, while those from NEMOf ^54^ were used in our initial split-FP designs. A total of six initial constructs were cloned and tested in Human Embryonic Kidney (HEK) 293T cells. While none of the variants responded to Ca^2+^, two of the designs exhibited strong and uniformly cytosolic expression (**Fig. 1b**). The two constructs, which were based on cpmScarlet3- and cpmScarlet-I3, were therefore selected for library screening with the goal of improving Ca^2+^ responses.

### Development of red fluorescent calcium indicators based on mScarlet fluorophores

To generate the first GECI library (Library 1), we varied the sequence identities of the flexible peptide linkers connecting the cpFP with the RS20 and CaM domains (**Fig. 2a**), as the linker sequence is known to tune biosensor responses.^2–4,38,64–66^ Overall, our library design strategy prioritized preservation of the native fluorophore, while allowing fine-tuned sampling of Ca^2+^-responsive sequences. Library 1 also sampled charged and hydrophobic residues in the first (denoted here as F1) and fourth (F4) positions of the cpFP, which immediately follow the first linker sequence. The F4 position corresponds to an arginine (R149) in mScarlet3^36,39^ and a methionine in mRuby.^37^ In the RCaMP crystal structure, the surface-exposed methionine (Met64 following RCaMP numbering) from mRuby interacts with three nonpolar residues, Pro360, Leu363 and Ile364, in the fourth alpha helix of CaM (**Supp. Fig. 1**).^3^ We therefore sampled multiple amino acid identities in this position. The final library comprised 6,144 linker variants and was used to populate the cpmScarlet3 and cpmScarlet-I3-based libraries, which were screened in parallel.

**Figure 2.**
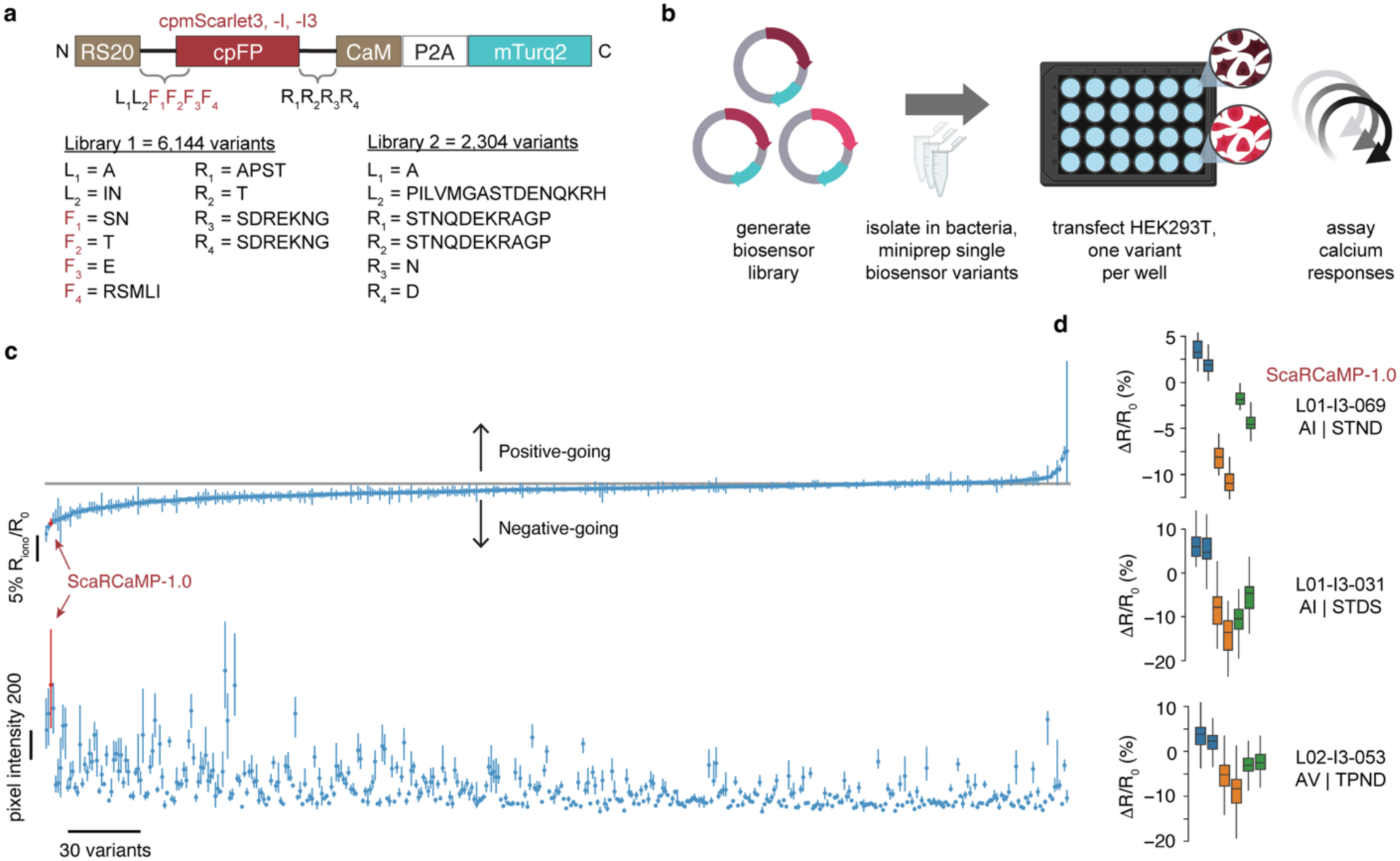
Development of mScarlet-based red fluorescent calcium indicators. (**a**) Schematic depicting cpFP-based GECI designs and linker variants sampled in Libraries 1 and 2. Library 1 tested variants using cpmScarlet3 or cpmScarlet-I3 as the cpFP, while Library 2 tested cpmScarlet-I3 or cpmScarlet-I as the cpFP. (**b**) Library screening workflow. HEK293T cells were transfected with individual library variants in multiwell plates and assayed for calcium responses using epifluorescence imaging. Created using BioRender.com. (**c**) Rank sorted average red-to-green ratio changes (percent change ionomycin response relative to baseline and EGTA; median ± 95% bootstrap confidence interval) for each of the 414 variants screened. Below, baseline brightness calculated as average pixel intensities are presented in the same order. Variant L01-I3-069, renamed ScaRCaMP-1.0, is indicated with a red arrow. (**d**) Fluorescence changes of selected mScarlet-I3-based sensor variants under three sequentially imaged conditions: baseline (blue), high calcium (orange; 10 μM ionomycin, 0.1 mM CaCl_2_), and low calcium (green; 2 mM EGTA added sequentially to the well). Each phase was imaged twice. Box plots quantify the average change in the ratio of red GECI fluorescence to green mTurquoise2 fluorescence (ΔR/R_0_), where R_0_ was calculated as the median across baseline and EGTA phases.

To perform library screening, HEK293T cells were seeded in multiwell glass-bottom plates, transfected with individual GECI variants, and imaged for Ca^2+^ responses in three phases (**Fig. 2b**). First, HEK293T cells were imaged in buffer to establish a baseline. Second, the cells were treated with 0.1 mM Ca^2+^ in the presence of 10 *μ*M ionomycin, an ionophore that shuttles Ca^2+^ across biological membranes. Third, Ca^2+^ was chelated with 2 mM EGTA. To control for motion artifacts, a green fluorescent protein (mTurquoise2) was co-expressed with the red GECIs, encoded on the same expression vector and separated by a P2A self-cleaving peptide sequence^67^ (**Fig. 2a**). Red fluorescence intensity was normalized against the green fluorescence signal within each region of interest (ROI), and calcium responses were plotted as the absolute change in red / green ratio (ΔR/R_0_; **Supp. Fig. 2**).

Of the variants screened, many of the cpmScarlet-I3 variants displayed a decrease in fluorescence intensity in response to Ca^2+^. Conversely, no Ca^2+^-responsive variants were observed in the cpmScarlet3 library. Four of the nonresponsive cpmScarlet3 variants were selected at random and sequenced, confirming construct viability and the absence of any unintended mutations, such as frameshift mutations (**Supp. Table 1**). The top-performing cpmScarlet-I3 variant showed a 13% decrease in normalized fluorescence in response to Ca^2+^ (**Fig. 2c, d**). While other variants showed larger response sizes (up to 20% fluorescence decreases), we found that L01-I3-069 had consistently low variance and high reproducibility across multiple experiments. This variant (L01-I3-069) was therefore renamed ScaRCaMP-1.0.

ScaRCaMP-1.0 contains a R149L mutation, and the left and right linker sequences are AI and STND, respectively (**Supp. Note 1**). These linker sequences are similar to those in jRCaMP1a (AI-PTDS). We also identified a variant (L01-I3-031) with linkers (AI-STDS) more similar to jRCaMP1a, and other variants sampling a Pro in the R1 linker position (L01-I3-032, AI-PTNK; L01-I3-028, AI-PTRK). All variants exhibited a fluorescence decrease in response to Ca^2+^ (**Fig. 2d; Supp. Table 1**), which contrasts with the large Ca^2+^-induced fluorescence increase observed in jRCaMP1a. This suggests that the mechanism driving fluorescence changes in cpmScarlet-I3 may differ from that in cpmRuby, the fluorophore used to construct jRCaMP1a.

A second library (Library 2) was designed around ScaRCaMP-1.0, sampling additional amino acid identities in the left and right linkers (**Fig. 2a**). Additional Ca^2+^ responsive variants were identified (**Fig. 2c, d; Supp. Table 1**), but they did not improve upon ScaRCaMP-1.0. In parallel, a version of Library 2 was prepared using cpmScarlet-I as the fluorophore. As with the cpmScarlet-I3-based library, the majority of biosensor variants expressed well in cells, and many responded to Ca^2+^. At the completion of library screening, a total of 414 variants had been screened. We then proceeded to further characterize ScaRCaMP-1.0.

### Characterization of ScaRCaMP-1.0

ScaRCaMP-1.0 protein was expressed in bacteria and purified for photophysical characterization. Excitation and emission spectra closely matched the spectral properties of the underlying fluorophore, mScarlet-I3.^36^ Addition of Ca^2+^ reduced fluorescence emission by 21% (**Fig. 3a**), similar to what was observed in cells. We also measured quantum yield and extinction coefficient for ScaRCaMP-1.0 in the presence and absence of Ca^2+^. Photophysical parameters are summarized in **Supp. Table 2**.

**Figure 3.**
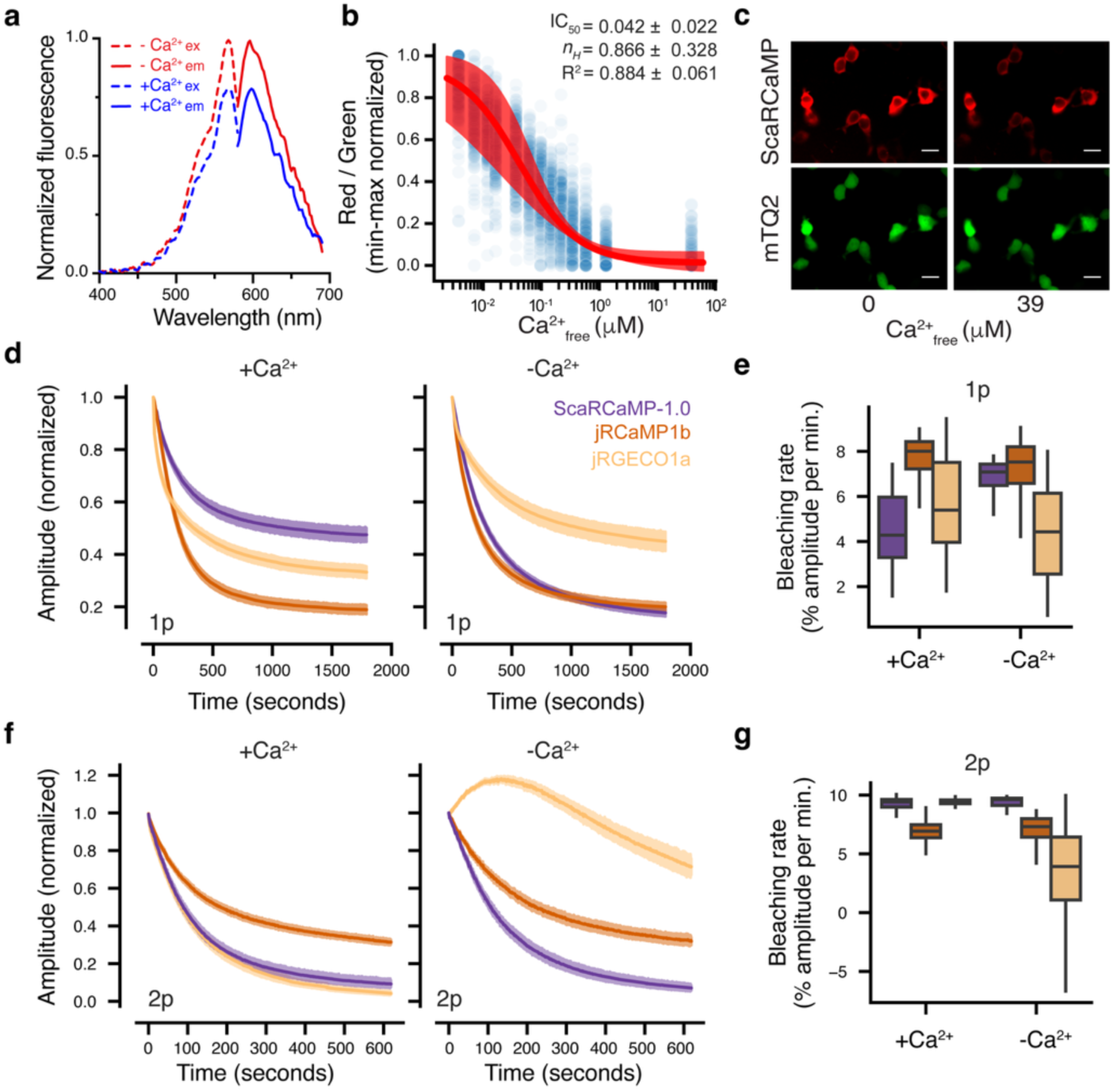
Photophysical properties of ScaRCaMP-1.0. **(a)** Excitation and emission spectra of purified ScaRCaMP-1.0 in Ca²⁺-free (-Ca^2+^, red) and Ca²⁺-saturated (+Ca^2+^, blue) conditions. **(b)** Ca²⁺ titration curve of ScaRCaMP-1.0 in permeabilized HEK293T cells. To account for motion artifacts and protein leak following permeabilization with digitonin, ScaRCaMP-1.0 (red) was normalized to mTurquoise2 (mTQ2, green) fluorescence, delivered via a bicistronic vector. N = 270 separate ROIs (blue dots) across 6 separate imaging sessions. Data from each session were fit with the Hill equation. Estimated dissociation constant for Ca^2+^ calculated as half-maximal inhibitory concentration (IC50) in μM, Hill coefficient (*n_H_*), and R^2^ values are presented as average ± standard deviation across the 6 sessions. Red line and shading represent average ± SD of fits across wells. **(c)** Representative changes in ScaRCaMP-1.0 fluorescence (top) in digitonin permeabilized HEK293T cells relative to mTurquoise2 fluorescence (bottom) in either 0 or 39 μM free Ca^2+^ buffer. Scale bar represents 20 μm. (**d**) 1-photon photobleaching curves in the presence (10 μM ionomycin, 1 mM Ca^2+^) and absence (BAPTA-AM and EGTA-treated) of Ca²⁺. Cells were imaged continuously under 555 nm, 52 mW illumination. (**e**) Quantification of 1-photon photobleaching rates per ROI, presented as the percent change in ΔF/F_0_ per minute. (**f**) 2-photon photobleaching curves with and without Ca^2+^, imaged continuously under 1050 nm, 98 mW illumination. (**g**) Quantification of 2-photon photobleaching rates per ROI, presented as in (e). Average, shading indicates 95% CI. 1p, +Ca^2+^: N_jRCaMP1b_ = 177, N_jRGECO1a_ = 175, N_ScaRCaMP-1.0_ = 168; 1p, -Ca^2+^: N_jRCaMP1b_ = 220, N_jRGECO1a_ = 133, N_ScaRCaMP-1.0_ = 145; 2p, +Ca^2+^: N_jRCaMP1b_ = 263, N_jRGECO1a_ = 359, N_ScaRCaMP-1.0_ = 251; 2p, -Ca^2+^: N_jRCaMP1b_ = 196, N_jRGECO1a_ = 327, N_ScaRCaMP-1.0_ = 188. To estimate bleaching curves, we took the first 10 minutes of fluorescence data for each ROI, then LOESS smoothed and computed the average gradient. For boxplots, boxes extend from the first quartile to the third quartile of the data, with a line specifying the median. The whiskers extend to the farthest datapoint within 1.5-fold of the interquartile range of the data. Outliers beyond these points are not plotted.

We next collected calcium titration curves in permeabilized HEK293T cells (**Fig. 3b, c**) as performed previously with other GECIs.^68^ GECI performance is known to differ *in vitro* versus in cells.^68–70^ It is therefore important to evaluate GECI responses in a cellular context. In cells, ScaRCaMP-1.0 showed a high apparent affinity for Ca^2+^ (IC_50_ = 42 ± 22 nM) and displayed low cooperativity (*n*_H_ = 0.87 ± 0.33), producing a linear response across a broad range of calcium concentrations. The majority of GCaMP- or RCaMP-derived GECIs exhibit high cooperativity with Hill coefficients nearing or exceeding 3.^2–4,38^ Low cooperativity GECIs, which are useful for monitoring graded Ca^2+^ responses, have also been developed, including those with high affinity (**Supp. Table 3**).^48,55,59,71–73^

Next, we assessed photostability during 1p and 2p imaging in intact mammalian cells. As bleaching rates can differ in the presence of Ca^2+^,^38^ we recorded photobleaching curves in cells treated either with ionomycin and 1 mM Ca^2+^ or with Ca^2+^ chelators (BAPTA-AM and EGTA; **Fig. 3d, f**). Consistent with previously published data, under 1p or 2p illumination, jRGECO1a resisted photobleaching in the absence of Ca^2+^ (**Fig. 3d-g**).^38^

Conversely, jRCaMP1b bleaching rates were similar in the presence and absence of Ca^2+^. ScaRCaMP-1.0 photobleaching was also unaffected by Ca^2+^ under 2p illumination (**Fig. 3f, g**) but was Ca^2+^-dependent under 1p illumination. Notably, under 1p illumination in the presence of Ca^2+^, ScaRCaMP-1.0 was less susceptible to photobleaching than either jRCaMP1b or jRGECO1a (**Fig. 3d, e**).

### ScaRCaMP-1.0 is insensitive to high-powered blue light and is compatible with optogenetic tools

We next assessed the compatibility of ScaRCaMP-1.0 with optogenetic stimulation and compared its performance against existing red GECIs (jRGECO1a, jRCaMP1a, jRCaMP1b, and PinkyCaMP). JRGECO1a, jRCaMP1a, and jRCaMP1b are widely used in *in vivo* calcium imaging experiments. PinkyCaMP is a recently described red GECI based on a modified cpmScarlet fluorophore that is reported to have no photoswitching behavior.^42^ We therefore included PinkyCaMP among our list of benchmarking GECIs. HEK293T cells were transfected with each of the five red GECIs, and co-transfected with the opsin CoChR^11,21^, which fluxes Ca^2+^ and other cations when activated by a pulse of blue light (**Fig. 4a-b**). When CoChR was present, photostimulation increased intracellular Ca^2+^ (**Fig. 4c**). Cells lacking CoChR were included as a control to measure red GECI responses to blue light only (**Fig. 4c, d**).

**Figure 4.**
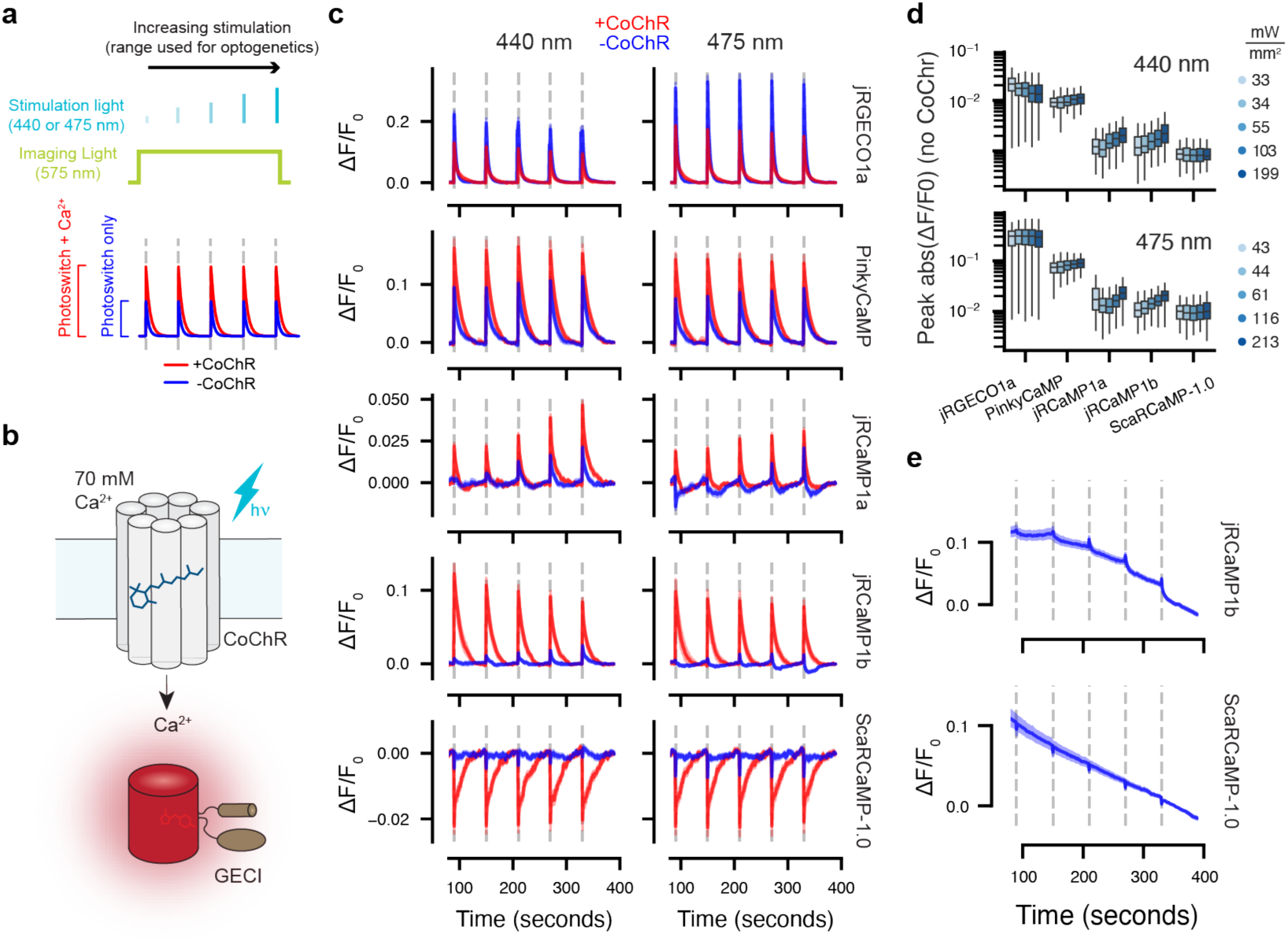
ScaRCaMP-1.0 reliably reports on calcium transients in optogenetics experiments in cultured cells. (**a**) Schematic of optogenetic stimulation protocol (440 nm and 475 nm, five sequential pulses of exponentially increasing power, with constant 575 nm illumination) and expected GECI responses. (**b**) Red GECIs (jRGECO1a, PinkyCaMP, jRCaMP1a, jRCaMP1b, and ScaRCaMP-1.0) were each co-expressed with CoChR, a light sensitive channel that fluxes cations (including Ca^2+^) in response to blue light. High extracellular Ca^2+^ (70 mM) was maintained to drive Ca^2+^ influx following channel opening. ATR added to increase CoChR photosensitivity. (**c**) Fluorescence intensity traces of red GECIs with and without CoChR under 440 nm or 475 nm stimulation, 5 second pulses. Shading 95% CI. 440 nm, +CoChR: N_jRGECO1a_ = 110, N_PinkyCaMP_ = 21, N_jRCaMP1a_ = 69, N_jRCaMP1b_ = 73, N_ScaRCaMP-1.0_ = 86; 440 nm, -CoChR: N_jRGECO1a_ = 249, N_PinkyCaMP_ = 77, N_jRCaMP1a_ = 200, N_jRCaMP1b_ = 298, N_ScaRCaMP-1.0_ = 201; 475 nm, +CoChR: N_jRGECO1a_ = 90, N_PinkyCaMP_ = 18, N_jRCaMP1a_ = 106, N_jRCaMP1b_ = 54, N_ScaRCaMP-1.0_ = 102; 475 nm, -CoChR: N_jRGECO1a_ = 242, N_PinkyCaMP_ = 125, N_jRCaMP1a_ = 231, N_jRCaMP1b_ = 324, N_ScaRCaMP-1.0_ = 241. (**d**) Quantification of peak blue light responses in (c), in the absence of CoChR. Data points are colored by illumination power according to the legend. (**e**) Indicated traces in (c), in the absence of CoChR. Here, a constant baseline was calculated across the whole session.

Channelrhodopsins can be activated by a broad spectrum of blue and cyan light.^12,74–76^ We therefore tested photoactivation in response to multiple wavelengths (either 440 nm or 475 nm light). In addition, during *in vivo* experimentation, the illumination power necessary for eliciting behavioral responses can vary. In particular, when light is introduced via a narrow fiber as done during fiber photometry experiments, peak irradiations spanning 10–200 mW/mm^2^ can be required.^77–82^ Few red fluorescent biosensors have been tested against power densities higher than 1 mW/mm^2^.^25–28,30,33,42,83^ We therefore tested a range of blue light power by increasing illumination power over a series of five pulses, ranging from 1 to 16% of illumination power, which corresponds to 33 to 199 mW/mm^2^ at 440 nm or 43 to 213 mW/mm^2^ at 475 nm in the imaging plane. Each pulse lasted either 1 or 5 seconds. Fluorescence was continuously recorded at 575 nm. Imaging began with a 90-second pre-stimulation baseline, stimulation pulse, and a 60-second post stimulation period between pairs of pulses (**Fig. 4a**). High Ca^2+^ (70 mM) was added to the external bath solution, consistent with previous experimental protocols for CoChR,^11^ to consistently drive maximal Ca^2+^ influx (**Fig. 4b**). In the absence of CoChR, responses to blue light alone could be measured (**Fig. 4c-e**).

At all wavelengths, pulse durations, and power levels tested, ScaRCaMP-1.0 consistently exhibited the smallest blue-light-induced fluorescence changes (peak ΔF/F_0, ScaRCaMP-1.0_ of 1 ± 0.3% and 1 ± 0.6% with 1 or 5 sec pulses of 475 nm light at 213 mW/mm^2^; **Fig. 4c-e, Supp. Fig. 3-4, Supp. Table 4-5**), with rapid kinetics that were notably faster than Ca^2+^ response times (**Fig. 4c, e; Supp. Fig. 3; Supp. Table 5**).

Consistent with previous findings, jRGECO1a, which is based on the mApple fluorophore, demonstrated the largest responses to blue light, approximately 30-fold larger than ScaRCaMP-1.0 (**Fig. 4c-e, Supp. Fig. 3-4**). For jRGECO1a, photoactivation is more pronounced in the absence of Ca^2+^ (**Fig. 4c**), which is consistent with previously reported data.^33,44^ This results in a regime where CoChR-induced Ca^2+^ influx counterintuitively reduces the apparent GECI response, and further emphasizes the incompatibility of jRGECO1a with optogenetics experiments.

JRCaMP1a and jRCaMP1b, which are based on the mRuby fluorophore, were not originally reported to exhibit photoswitching behavior in response to lower powered blue light.^38^ Still, at high power we observed clear photoswitching, approximately twice that of peak ScaRCaMP-1.0 photoactivation (**Fig. 4d, Supp. Fig. 3-4**). This is consistent with the photochromic behaviors observed in the mRuby lineage of fluorescent proteins.^36,37^ We also noted that high powered pulses of blue light produced complex baseline shifts in the jRCaMP1b signal that were absent in ScaRCaMP-1.0 (**Fig. 4e, Supp. Fig. 3**). Baselines exhibiting multiexponential decay can pose a challenge for baseline correction, giving rise to artifacts such as the subtle but noticeable “undershoots” evident in the baseline-corrected traces for jRCaMP1b following 475 nm light exposure (**Fig. 4c, e**). Conversely, the monoexponential decay of ScaRCaMP-1.0 traces lends itself to straightforward baseline correction (**Fig. 4e, Supp. Fig. 3**).

Surprisingly, in our hands PinkyCaMP did not demonstrate blue-light photostability, even though it is derived from cpmScarlet and has been reported to lack photoswitching behavior.^42^ Under our experimental conditions, PinkyCaMP blue light responses exceeded those of all red GECIs tested except for jRGECO1a (**Fig. 4c, Supp. Fig. 3-4, Supp. Tables 4-5**), with peak responses nearly 5-fold greater than that of ScaRCaMP-1.0. The kinetics of the PinkyCaMP photoswitching response also differed from that of ScaRCaMP-1.0 (**Fig. 4c, Supp. Fig. 3**). Following a blue light pulse, ScaRCaMP-1.0 rapidly returned to baseline, whereas PinkyCaMP displayed slower off-rates that mimicked the kinetics of a Ca^2+^ response (**Fig. 4c**). We speculate that some or all of the 11 mutations introduced into the PinkyCaMP fluorescent protein during the course of biosensor screening^42^ may be important for determining photoswitching behavior. The cpmScarlet-I3 fluorophore is unmodified in ScaRCaMP-1.0, and the high degree of blue-light photostability observed in the native fluorophore^36^ seems to have been largely preserved in ScaRCaMP-1.0. Thus, ScaRCaMP-1.0 accurately reports intracellular Ca^2+^ changes induced by blue-light photoactivation of CoChR.

### ScaRCaMP-1.0 reports on neural activity in awake behaving animals

To confirm the compatibility of ScaRCaMP-1.0 with *in vivo* imaging, we conducted dual-color fiber photometry in the striatum of awake head-fixed mice to test its performance with existing green and red GECIs. In the first phase of *in vivo* testing, we co-injected experimental mice with a mixture of ScaRCaMP-1.0 (AAV9-CAG-ScaRCaMP-1.0) and jGCaMP8m (AAV9-Syn-jGCaMP8m-WPRE) in the striatum, followed by implantation of fiber optic cannulae. Photometric recordings were conducted for both green and red emission channels 7–51 days post-implantation (**Fig. 5a, b**) while head-fixed mice ran on a spinning disk. We found that ScaRCaMP-1.0 and jGCaMP8m displayed distinct traces from the isosbestic signal (**Fig. 5c**), validating the ability of ScaRCaMP-1.0 to detect *in vivo* Ca^2+^ transients.

**Figure 5.**
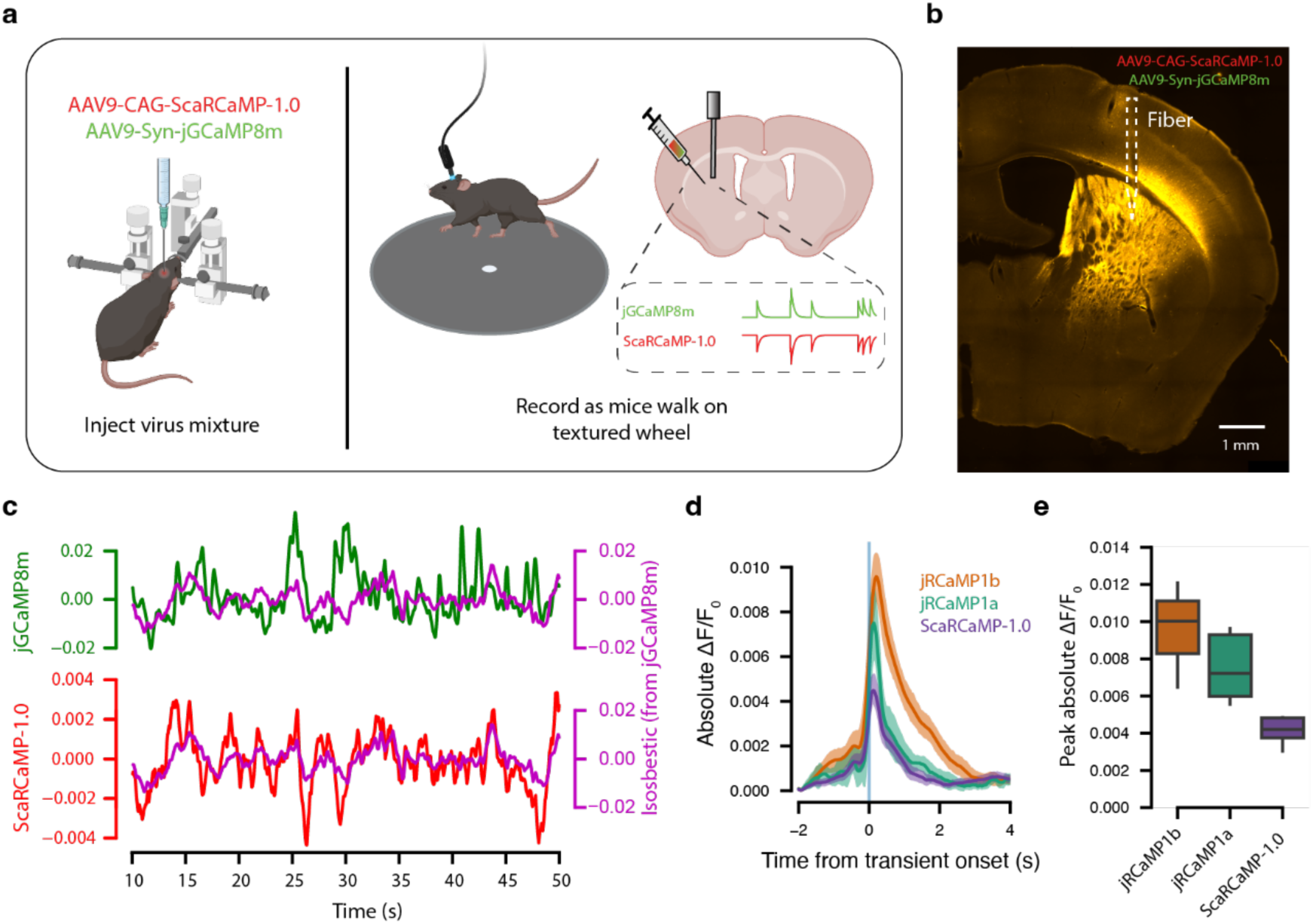
ScaRCaMP-1.0 reports on neural activity in awake behaving animals. (**a**) Schematic of *in vivo* experimental paradigm created using BioRender.com. (**b**) Representative histological section showing viral expression (ScaRCaMP-1.0: jGCaMP8m, 5:1) in the DLS (scale bar: 1 μm). (**c**) Representative fiber photometry traces from a mouse co-expressing jGCaMP8m and ScaRCaMP-1.0, showing concurrent Ca^2+^-dependent fluorescence dynamics. Magenta purple traces represent the isosbestic signal from jGCaMP8m excited at 405 nm. (**d**) Calcium responses comparing red GECI signals. Here we took 6 second windows surrounding threshold crossing 2 SD above the mean (jRCaMP1b and jRCaMP1a) or below the mean (ScaRCaMP-1.0). Shown is average, with shading indicating 95% CI. (**e**) Absolute fluorescence peaks for ScaRCaMP-1.0, jRCaMP1b, and jRCaMP1a.

In the second phase of *in vivo* testing, we directly compared ScaRCaMP-1.0 to two widely used red GECIs, jRCaMP1a and jRCaMP1b, focusing on kinetic properties and peak fluorescence responses. Either AAV9-Syn-NES-jRCaMP1a or AAV9-Syn-NES-jRCaMP1b was co-injected with jGCaMP8m into the same striatal region and recorded following the previous protocol for 7–34 days. Consistent with previous reports, jRCaMP1b showed the largest responses, followed by jRCaMP1a, with ScaRCaMP-1.0 producing smaller but clearly detectable signals (**Fig. 5d, e**). This establishes that ScaRCaMP-1.0 can be used to detect intracellular Ca^2+^ changes *in vivo*.

### Mutation of surface-exposed lysines on mScarlet-I3 alters calcium responses

To understand the potential mechanism underlying the fluorescence change in ScaRCaMP-1.0, we turned to AlphaFold3.^43^ Historically, it has been challenging to gain structural information on both the holo (Ca^2+^-bound) and apo (Ca^2+^-free) states of GECIs.^84^ We therefore reasoned that structural predictions may be useful for hypothesis generation.

We submitted the ScaRCaMP-1.0 primary amino acid sequence to the AlphaFold3 server, without modeling Ca^2+^ ions, to obtain 40 structures across 8 seed groups. When we overlaid the structures, we found that AlphaFold3 predictions sampled multiple states (**Supp. Fig. 5**), including conformations that could be consistent with an “open” state or with a “closed” state (**Fig. 6a, Supp. Fig. 6**). Upon closer inspection, we found suggestion of a potential mechanism for the fluorescence changes in ScaRCaMP-1.0. In the predicted “closed” state, Ile79, which resides in the left linker, interacts with a surface-exposed lysine residue (Lys132), sealing the beta-barrel (**Fig. 6a**). In the predicted “open” state, Ile79 switches to a second surface-exposed lysine residue (Lys96) on the opposite side of the beta-bulge, opening a gap in the beta-barrel. Such a lesion may reduce fluorescence intensity by allowing entry for water molecules which could reduce fluorescence via collisional quenching^85–87^ or other means.^36,39,88^ We therefore reasoned that the “open” state may represent the low-fluorescence state, while the “closed” state may represent the high-fluorescence state. AlphaFold3 structural predictions did not suggest a mechanistic role for the R149L mutation.

**Figure 6.**
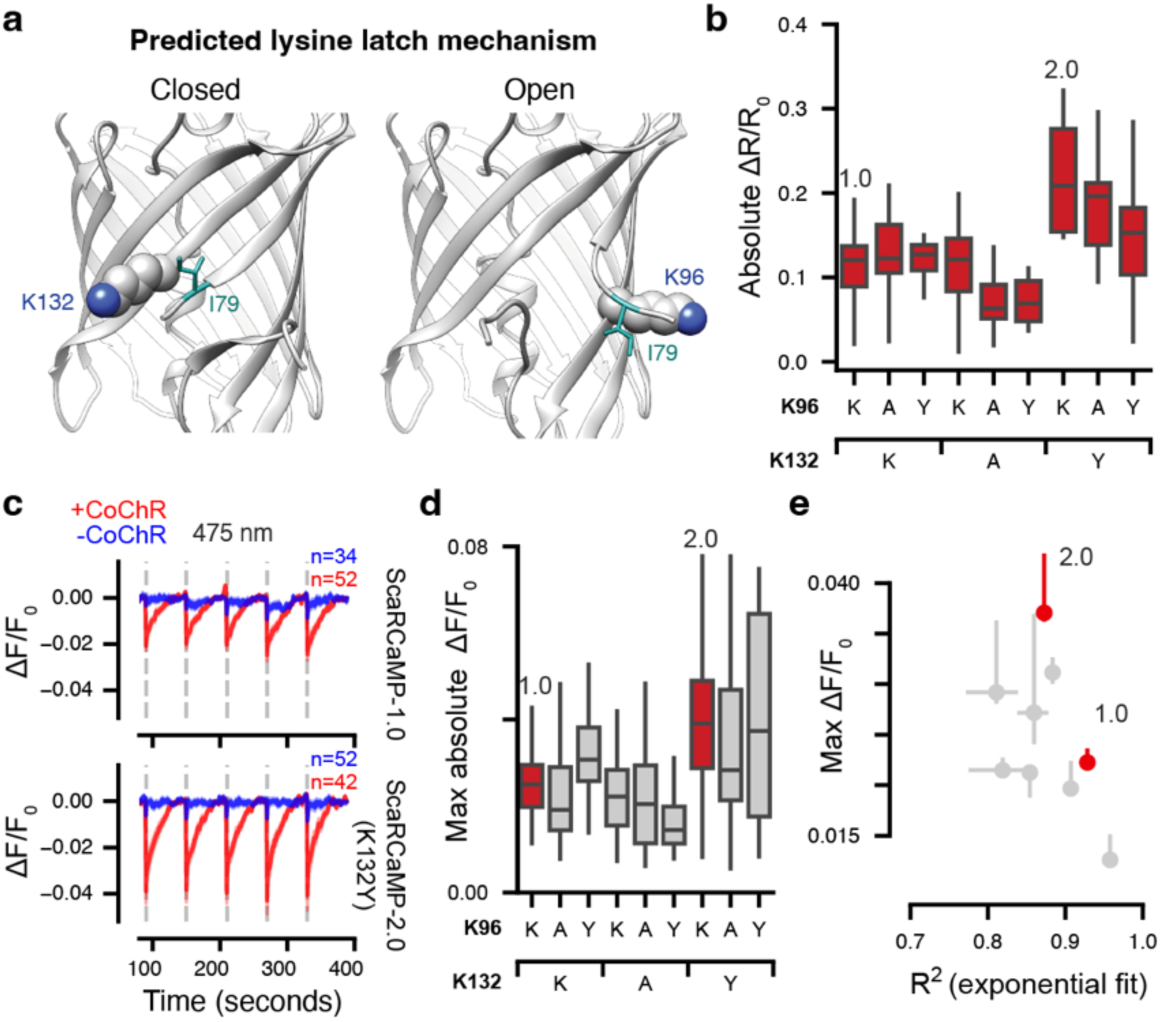
Mechanism underlying ScaRCaMP-1.0 calcium responses and the development of ScaRCaMP-2.0. **(a)** AlphaFold3 structural predictions for ScaRCaMP-1.0 suggest a lysine latch on the fluorescent protein surface. Cartoon rendered in UCSF Chimera v1.19.^97^ **(b)** Ca^2+^ responses in HEK293T cells expressing single and double mutants. Absolute ΔR/R_0_ is the change in the red-to-green ratio (normalized to mTurquoise2), when cells receive Ca^2+^ + ionomycin, relative to EGTA. Cells were pre-depleted of Ca^2+^ with EGTA treatment prior to imaging. **(c)** Optogenetically (CoChR) induced Ca^2+^ responses as in Fig. 4, 475 nm, 5 second pulse. **(d)** Quantification of responses from **(c)**. **(e)** Ca^2+^ responses (ΔF/F_0_) in the presence of CoChR, plotted as a function of blue-light photoswitching in the absence of CoChR. Here photoswitching is quantified by goodness of fit (R^2^) to a monoexponential function, where R^2^=1 is a perfect fit. **, p<0.01, ***, p<0.001, Mann-Whitney U test compared with ScaRCaMP-1.0. Error bars indicate 95% bootstrap confidence interval.

We were intrigued by the suggestion that the two “lysine latches” residing on the surface of mScarlet-I3 may be important for the calcium responses in ScaRCaMP-1.0. Recently, crystal structures of other fluorescent biosensors have pointed to the existence of surface latches that are critical for biosensor function.^89^ We therefore decided to explore this hypothesis further.

We tested the validity of the structural predictions using site-directed mutagenesis. We generated eight mutants: 4 single point mutants (K96A, K96Y, K132A, and K132Y) intended to either strengthen (Y) or weaken (A) latch interactions, and 4 double mutants (AA, AY, YA, and YY, for positions 96 and 132 respectively). Mutants were tested first by treating HEK293T cells with ionomycin and Ca^2+^ or EGTA (**Fig. 6b**), and later by optogenetically increasing intracellular Ca^2+^ via CoChR activation (**Fig. 6c-d**). Strikingly, a single point mutant, K132Y, increased Ca^2+^ responses without compromising the blue light resistance of the sensor (**Fig. 6e**, **Supp. Table 6**). We renamed this variant ScaRCaMP-2.0. ScaRCaMP-2.0 displays average ΔF/F = -22% (**Fig. 6b**), establishing feasibility for increasing signal size while maintaining blue light compatibility. K132Y double mutants also showed increased Ca^2+^ responses, while K132A mutants showed reduced Ca^2+^ responses (**Fig. 6b, d**), supporting the hypothesis that K132 is important for ScaRCaMP responses.

## Discussion

In this study, we present data showing that mScarlet-derived fluorophores can be used to develop bright and photostable Ca^2+^ indicators. We report two new red fluorescent GECIs, called ScaRCaMP-1.0 and ScaRCaMP-2.0. While their Ca^2+^ response sizes (ΔF/F_0_ = - 13% and -22% for ScaRCaMP-1.0 and -2.0, respectively) are more modest compared with existing red GECIs, ScaRCaMP GECIs demonstrate exceptional photostability, making them particularly well-suited for optogenetic regimes in which blue-light tolerance may be prioritized over response size. Overall, the results presented here address limitations related to blue-light-induced photoactivation of red GECIs and offer an alternative design strategy for producing optogenetics-compatible tools.

In our approach, we first systematically tested two GECI design topologies (cpFP and split-FP) with each of three red fluorescent proteins with desired fluorescence properties (mScarlet3, mScarlet-I3, and mCherry-XL). We then subjected the highest expressing designs (cpmScarlet3 and cpmScarlet-I3) to library screening and identified linker sequences that yield Ca^2+^ responses. We also incorporated cpmScarlet-I into later library screens. The cpmScarlet-I3 and cpmScarlet-I fluorophores readily yielded Ca^2+^- responsive variants that decrease their fluorescence in response to Ca^2+^. It is not yet clear if the inverted response is inherent to these two fluorophores, or if it is a product of the linker sequences sampled by the library. Inverse-response GECIs are rare but have been useful in monitoring Ca^2+^ changes in worms, imaging baseline states, multiplexing with positive-going sensors, and providing reliable signals during miniscope implantation in anesthetized mice.^59,71^

We named the lead library variant ScaRCaMP-1.0, which is based on the cpmScarlet-I3 fluorophore. Most importantly, ScaRCaMP-1.0 is insensitive to blue light, showing minimal (1%) fluorescence changes in response to intense (>200 mW/mm^2^) pulses of 475 nm light, spanning the highest power ranges used in *in vivo* optogenetics experiments. Comparatively, peak blue light responses from ScaRCaMP-1.0 were roughly 30-fold smaller than that of jRGECO1a and 2-fold smaller than that of jRCaMP1a and jRCaMP1b. We also compared ScaRCaMP-1.0 to another recently developed cpmScarlet-based GECI, called PinkyCaMP.^42^ Despite being derived from cpmScarlet, under the conditions tested here PinkyCaMP showed clear photoswitching behavior, which may arise from a subset of mutations introduced during screening, most likely those modifying inward-facing residues neighboring the chromophore. In designing ScaRCaMP-1.0, mutations were restricted to regions within and immediately adjacent to the peptide linkers to minimize the likelihood of rescinding the beneficial properties of the fluorophore. Consequently, the resistance to photoactivation imparted by cpmScarlet-I3 was largely preserved in ScaRCaMP-1.0. These observations further emphasize the importance of maintaining the native chromophore environment of mScarlet FPs, as it may be critical for preserving blue-light photostability.

As a demonstration of its *in vivo* performance, we expressed ScaRCaMP-1.0 in the striatum. ScaRCaMP-1.0 yielded signals detectable by fiber photometry in awake head-fixed mice, with response sizes approaching those of jRCaMP1a and kinetics that were faster than jRCaMP1b but slower than jGCaMP8m. With future variants of ScaRCaMP-1.0, improved *in vivo* signals may be achieved. Still, our sensor fits within specific *in vivo* imaging niches. Our experiments using jGCaMP8m demonstrate the use of blue-light photostable red GECIs like ScaRCaMP-1.0 in multiplex imaging with green fluorescent indicators. And in cases where optogenetic stimulation requires high powered blue light, ScaRCaMP-1.0 may prove to be a preferable tool relative to other red GECIs.

We also explored the mechanism underlying ScaRCaMP-1.0 calcium responses. Guided by AlphaFold3 structural predictions, we identified two lysines on the surface of mScarlet-I3 that were potentially involved in ScaRCaMP’s Ca^2+^ responses. We hypothesized that the isoleucine within the linker region may alternately interact with each lysine residue to induce conformational changes in the FP. We found that mutation of each lysine, alone or in concert, could alter ScaRCaMP response sizes, and that a single point mutant, K132Y, improved Ca^2+^ responses while maintaining blue-light photostability. This is notable as photoswitching and response size can be coupled (see PinkyCaMP, **Fig. 4c**). In addition, because both lysine residues are intrinsic to the FP, it is feasible that this aspect of the ScaRCaMP mechanism may be translated to develop other mScarlet-I3-based biosensors. Interestingly, both lysines are also conserved in mScarlet and mScarlet3, which failed to yield calcium responses in our screens. This suggests that the lysine latches, while important, are not sufficient for ScaRCaMP responses.

One final noteworthy feature of ScaRCaMP-1.0 is that it displays low cooperativity and a linear response to Ca^2+^. While sharp transitions between dark and bright states are suitable for spike detection, low cooperativity GECIs are useful for monitoring graded Ca^2+^ responses.^71^ Existing GECIs with Hill coefficients near 1 include high affinity variants of red cpFP GECIs (R-CaMP2, R-GECO2L),^48^ a cpFP GECI with a lifetime response (Tq-Ca-FLITS),^72^ split-FP GECIs (NCaMPs, iYTnC),^55,59^ and GECIs with an inverted response (Inverse-pericam, IP2.0, iYTnC) (**Supp Table 3**).^59,71^ While lower cooperativity is often accompanied by a reduction in dynamic range, certain GECIs have achieved very large response sizes with low Hill coefficients (NCaMP9, NCaMP10, IP2.0).^55,71^ It is therefore feasible that, in future ScaRCaMP variants, higher response sizes may not come at the expense of cooperativity.

There are limitations to ScaRCaMP-1.0 and ScaRCaMP-2.0. Future iterations would benefit from further improvements to Ca^2+^ response sizes. On average, jRCaMP1b has roughly 5-fold and 2-fold larger Ca^2+^ responses than ScaRCaMP-1.0 and ScaRCaMP-2.0, respectively, and jRCaMP1b may still be usable in optogenetic regimes calling for low powered blue light.^90^ ScaRCaMP may also benefit from faster response kinetics and reduced Ca^2+^ affinity. Additionally, we did observe that certain ScaRCaMP mutants demonstrated an increase in blue-light photoactivation. Thus, future screening paradigms will benefit from incorporating blue-light responsiveness as a selection criterion during screening. Likewise, structural investigation of the Ca^2+^-bound and -unbound states would deepen our mechanistic understanding of the coupling between photoswitching and response size. Finally, given the robust expression profile, resistance to blue-light-induced photoswitching, and functionality in a calcium indicator design, we anticipate that the mScarlet-I3 fluorophore may serve as a promising scaffold for the development of additional photostable red fluorescent biosensors.

## Materials and Methods

### Fluorescence Microscopy

Mammalian cells were imaged with a Nikon ECLIPSE Ti-2 widefield microscope equipped with a Lumencor Spectra III light engine and a Hamamatsu Orca Flash 3.0 CMOS sensor. Images for red (555 nm excitation, 52 mW power, 50 ms exposure time) and green (475 nm excitation, 21 mW power, 20 ms exposure time) fluorescence were collected with a Plan Apo λD 20X/0.8NA objective and passed through a Semrock filter cube (BrightLine® full-multiband filter set, LED-DA/FI/TR/Cy5/Cy7-A-000). Red emission was additionally collected through an AT605/52 nm bandpass filter (Chroma). Green emission was collected through an ET520/40 nm bandpass filter (Chroma). Images were analyzed using custom Python code. To correct for motion, we used the “rigid_body” setting for PyStackReg,^91^ with each frame using the previous frame as a reference. Then, for identifying regions of interest, we used the “cyto2” model in Cellpose^92^ to identify individual cells. Fluorescence intensity was then estimated per cell and per frame by taking the mean across all pixels in each region of interest.

Two-photon (2p) imaging was performed on a Bruker Ultima Investigator upright multiphoton microscope, with 2p excitation at 1050 nm (Coherent Chameleon Discovery, 80 MHz Ti:sapphire laser) focused through a water immersion Plan Fluorite objective (Nikon N16XLWD-PF, 16X/0.8 NA) and laterally scanned across the sample (512 x 512 pixels) with a 1.6 μs per pixel dwell time. Fluorescence emission was passed through a Longpass Dichroic Beamsplitter (Chroma, T565lpxr) followed by an emission filter (Chroma, ET595/50m) and collected onto a GaAsP detector (Hamamatsu H10770) connected to a Becker & Hickl SPC-150 time-correlated single-photon counting (TCSPC) module.

### Plasmid Design and Cloning

All PCR reactions were performed using Q5 Hot Start High-Fidelity 2X Master Mix (NEB), and Gibson Assembly reactions were performed using NEBuilder HiFi DNA Assembly (NEB). All plasmids were sequence verified with nanopore sequencing (Plasmidsaurus). All proteins carry an N-terminal His-tag.

Split-FP and cpFP GECIs were synthesized by Genewiz. Split-FP GECIs were cloned by Genewiz into a pcDNA3.1 backbone containing a nuclear exclusion sequence (NES). CpFP GECIs were cloned by Genewiz into a pDx backbone derived from pDx-mScarlet3 (Addgene #189754). Gibson Assembly was then used to introduce an NES sequence that was PCR amplified from pGP-CMV-NES-jRCaMP1b (Addgene #63136) using F’ and R’ primers (**Supp. Table 7**).

For protein expression in bacteria, ScaRCaMP-1.0 was subcloned into a pRSETB bacterial expression vector by PCR amplifying the ScaRCaMP-1.0 gene (pRSETB_NES_gib_F and pRSETB_CaM_gib_R) and inserting it via 2-fragment Gibson into a pRSETB backbone (pRSETB-KanR-LiLac, Addgene # 184569) digested with NdeI-HF (NEB) and BspEI (NEB).

For optogenetics experiments, pGP-CMV-CoChR-GFP was cloned by PCR amplifying the CoChR-GFP gene (CoChR_F and CoChR_R) from pAAV-Syn-CoChR-GFP (Addgene # 59070) and inserting it via 2-fragment Gibson into a pGP backbone digested with NheI-HF (NEB) and NotI-HF (NEB).

### Cell Culture and Transfection

HEK293T (ATCC, #CRL-3216) or HEK293FT (Invitrogen R70007) cells were cultured in 5% CO_2_ at 37°C. Cells were maintained in Dulbecco’s Modified Eagle’s Medium (DMEM, high glucose, pyruvate; Gibco) supplemented with 10% Fetal Bovine Serum (FBS, One Shot; Gibco) and 1% Penicillin-Streptomycin (PS, 10,000 U/mL; Gibco). Cells were subcultured when they reached approximately 80% confluency. For transfection, cells were seeded on 1 cm^2^ glass coverslips or in glass-bottom 12 or 24-well plates (Fisher Scientific Cellvis) pre-coated with a protamine solution (1 mg/mL in MilliQ water; Sigma-Aldrich) to promote adherence. The seeding medium consisted of FluoroBrite™ DMEM (Gibco) supplemented with 10% FBS, 1% PS, and 1X GlutaMAX™ (Gibco).

Transfections were performed 24–48 hours after seeding or when cells reached 40–60% confluency using Effectene Transfection Reagent (Qiagen) or CalPhos Mammalian Transfection Kit (Takara bio #631312). Each well received a total of 0.5 μg (glass coverslip) or 0.4 μg (12-well plate) or 0.3 μg (24-well plate) of plasmid DNA. Cells were incubated for an additional 24–48 hours before imaging. For experiments with CoChR, media was supplemented with 1 μM all-trans-retinal (ATR, Sigma). For optogenetic stimulation, wells were co-transfected with each red GECI and CoChR at a 3:1 ratio and subsequently incubated in ATR-containing high-Ca^2+^ imaging buffer.

### Library Generation

Library 1 and Library 2 were generated using degenerate oligonucleotides containing codon variation within and adjacent to the linker regions, flanked by an annealing sequence and a Gibson overhang sequence. Degenerate oligos were used as primers in a standard PCR reaction (Q5 Hot Start High-Fidelity 2X Master Mix, NEB) to amplify cpmScarlet3, cpmScarlet-I3, or cpmScarlet-I-containing templates synthesized by Genewiz. Primers used to populate Library 1 were: SI3_L01_F and SI3_L01_R. Primers for Library 2 used to amplify cpmScarlet-I3 and cpmScarlet-I, respectively, were: SI3_L02_F and SI3_L02_R; SI_L02_F and SI_L02_R (**Supp. Table 8**).

To prepare the vector backbone, we first generated staging plasmids containing the cpmScarlet3-, cpmScarlet-I3-, and cpmScarlet-I-based GECIs synthesized by Genewiz. For both libraries, the staging plasmids consisted of a pDx backbone, either with or without a P2A-mTurquoise2 sequence (synthesized by Genewiz) inserted via Gibson Assembly into a BsrGI site immediately following the GECI sequence. For Library 2, a second staging plasmid was prepared using a pGP backbone from pGP-CMV-NES-jRCaMP1b (Addgene #63136)^38^, with the P2A-mTurquoise2 sequence inserted, following the observation of plasmid dimers using the pDx backbone.

Plasmid libraries were constructed using 2-fragment Gibson assembly of one vector backbone and one insert (NEBuilder HiFi DNA Assembly, NEB). The pDx vectors were digested with SacI-HF (NEB) and SapI (NEB), and pGP vectors were linearized by PCR (pGP_F and pGP_R). Gibson products were used to transform XL-10 Gold Ultracompetent Cells (Agilent Technologies) following the manufacturer’s instructions. The recovered mixture was plated on LB-agar plates supplemented with 50–100 μg/mL Kanamycin and incubated overnight at 37°C. Individual colonies were selected, cultured and miniprepped using QIAprep Spin Miniprep Kit (Qiagen). Following screening, lead sensors were subcloned into a pAAV backbone from pAAV-CAG-GCaMP6f-WPRE-SV40 (Chen 2013; Addgene #100836)^4^.

### Library Screening

HEK293T or HEK293FT cells were transfected as described above, using individual minipreps from the plasmid DNA library. On the day of imaging, cells were transferred to pre-warmed Imaging Buffer (0.1 mM CaCl_2_, 10 mM HEPES, 145 mM NaCl, 2.5 mM KCl, 10 mM D(+)-glucose, 1 mM MgCl_2_, pH 7.4 at room temperature) 20 minutes prior to imaging. Imaging was performed in three phases: (1) baseline, in Imaging Buffer, (2) high Ca^2+^, where 10 μM ionomycin was added to the well, and (3) low Ca^2+^, where 2 mM EGTA was added to the well. For each phase, two images per channel were acquired within the same field of view. Red GECI signals were imaged using 555 nm excitation (50 ms exposure, 52 mW power), and green fluorescence from the co-expressed mTurquoise2 was excited with 475 nm excitation (20 ms exposure, 21 mW power). Red and green emission channels were collected in rapid series. Max calcium response (ΔF/F_0_) is defined as the normalized change in fluorescence ((min[ionomycin]-max[baseline])/max[baseline]). Calcium responses are also calculated as a ratio of red to green channels (ΔR/R_0_), defined as the average change in the ratio of red GECI fluorescence to green mTurquoise2 fluorescence, where R_0_ was calculated as the median across baseline and EGTA phases. Data were analyzed using the same pipeline described in “Fluorescence Microscopy.”

### Protein Purification

BL21(DE3) Competent *E. coli* (NEB) were transformed with pRSETB-ScaRCaMP-1.0, which carries an N-terminal His-tag. 100 ml LB broth with 100 μg/mL Kanamycin was inoculated with 5 mL of overnight culture and grown at 37°C shaking overnight at 240 rpm. Protein expression was induced with a final concentration of 1 mM IPTG at room temperature shaking at 240 rpm for 48 hours. Cell pellets were stored at -80°C.

For purification, cell pellets were thawed on ice, resuspended in lysis buffer (50 mM HEPES, 500 mM NaCl, 1 mM TCEP, pH7.5, 10% glycerol, Pierce Protease Inhibitor Mini Tablets (Thermo), DNAse (Millipore Sigma)) and sonicated on ice (10 seconds pulse on, 10 seconds pulse off) for a total time of 30 minutes. Lysates were clarified for 5–10 min at 6,500 g at 4–6 °C. HisPur Ni-NTA Resin (Thermo) was loaded into an Econo-Column (BioRad) for affinity purification. The column was washed and equilibrated with Wash Buffer (200 mM MOPS, pH 7.4, 500 mM KCl, 10 mM imidazole). Lysates were loaded onto the gravity column, washed with 10 column volumes of Wash Buffer, then eluted with three column volumes of Elution Buffer (50 mM MOPS, pH 7.4, 300 mM KCl, 500 mM imidazole). Eluted protein was dialyzed in Dialysis Buffer (25 mM Tris-HCl, pH 7.3, 90mM KCl, 10 mM NaCl) using a 10K MWCO Slide-A-Lyzer**^TM^** G3 Dialysis Cassette (Thermo) while stirring at 4 ° C overnight. Protein was concentrated using a Pierce™ Protein Concentrator (10K MWCO, Thermo).

### Photophysical Characterization

Fluorescence excitation and emission spectra were measured using a BioTek Synergy H4 Hybrid plate reader. ScaRCaMP-1.0 protein was diluted in 25 mM Tris-HCl, pH 7.3, 90mM KCl, 10 mM NaCl at room temperature with 1 mM EGTA or 1 mM CaCl_2_. Spectra were acquired in triplicate and then averaged. Excitation spectra were measured in 2 nm increments from 350–590 nm, measuring emission at 620 nm. Emission spectra were measured in 2 nm increments from 580–700 nm, with 560 nm excitation. Excitation and emission spectra were normalized and plotted with 2^nd^ order smoothing, 3 neighbors each side, in GraphPad Prism 10.

### Quantum Yield

To determine the relative quantum yield of ScaRCaMP-1.0, fluorescence measurements were made using a Cary Eclipse Fluorescence Spectrophotometer (Agilent) controlled by Cary Eclipse Scan Application (Agilent), and absorbance spectra were recorded on a Cary 3500 UV-Vis Compact (Agilent).^93,94^ Protein was measured in Zero or 39 μM Free Calcium Buffer from Calcium Calibration Buffer Kit #1 (Thermo) at room temperature in triplicates. Rhodamine 101 (Sigma-Aldrich) in 100% ethanol was used as a standard. Absorbance spectra were recorded from 200–800 nm with 1 nm interval, 2 nm spectral bandwidth, and 0.02 s averaging time. Fluorescence emission spectra were recorded from 545–700 nm (5 nm slit), using 540 nm excitation. Spectra were integrated and analyzed in GraphPad Prism 10.

### Extinction Coefficient

Extinction coefficients were determined as in Gadella et al 2023.^36^ Purified ScaRCaMP-1.0 protein was diluted in either Zero or 39 µM Free Calcium Buffer from the Calcium Calibration Buffer Kit #1 (Thermo). Absorbance spectra were measured on a Cary 3500 UV-Vis Compact (Agilent) across a wavelength range of 260–700 nm, with a step size of 1 nm. Buffer alone was used as a background reference. Minor offsets were corrected by subtracting the blank solution absorbance and the average absorbance value across the wavelength range 670–680 nm. The concentrations of ScaRCaMP-1.0 protein were determined by measuring the absorbance following alkaline denaturation and assuming ε = 44,000 M^-1^ cm^-1^ at 457 nm. Extinction coefficients of ScaRCaMP-1.0 were calculated by dividing the peak absorbance maximum by the concentration of protein according to Beer’s law.

### Calcium Titrations

Calcium titration curves were measured in permeabilized HEK293T cells. HEK293T cells were transfected with pGP-CMV-NES-ScaRCaMP-1.0-P2A-mTurquoise2. Digitonin (2–5 µM; Sigma-Aldrich) was added to both the Zero and the 39 µM Free Calcium Buffers from Calcium Calibration Buffer Kit #1 (Thermo). Prior to imaging, cells were washed twice with D-PBS (Gibco) and then exchanged into Zero Free Calcium Buffer with digitonin. Cells were allowed to permeabilize for 5 minutes prior to imaging. A reciprocal dilution method was followed to achieve 14 conditions with free Ca^2+^ ranging from 0 to 39 µM. Changes in ScaRCaMP-1.0 fluorescence were normalized against mTurquoise2 to account for motion artifacts and gradual protein leakage from permeabilized cells. Data were pre-processed using the same pipeline described in “Fluorescence Microscopy.” Then, data from all ROIs (cells) in a well were used to fit the parameters of a standard Hill curve. Data were fit using the “least_squares” function in scipy.optimize. Cells with >10% decrease in mTurquoise2 fluorescence between the first and final imaging datasets were excluded from analysis. Cells with an average pixel intensity of 200 or less, or with a fluorescence ratio change less than 10% or greater than 100% were also excluded from further analysis.

### Photobleaching

For photobleaching experiments, HEK293T cells were transfected with sensor plasmids (pGP-CMV-NES-jRCaMP1b, pGP-CMV-NES-jRGECO1a, or pGP-CMV-ScaRCaMP-1.0) and imaged 24–48 hours later. For 1p imaging, cells were seeded in glass-bottom multiwell plates. For 2p imaging, cells were seeded on glass coverslips. Glass was coated with protamine. Coverslips were cut to size and placed in an imaging chamber filled with Imaging Buffer supplemented with either (1) 1 mM CaCl_2_ and 10 µM ionomycin, or (2) 10 µM ionomycin, 2 mM EGTA, and 50 µM BAPTA-AM. Cells were imaged continuously under 1p excitation (555 nm, 52 mW) or 2p excitation (1050 nm, 98 mW). Data were analyzed using the same pipeline as described in “Fluorescence Microscopy” and plotted as percent normalized by initial pixel intensity over time per ROI.

### Photoswitching

Cells were co-transfected with CoChR (pGP-CMV-CoChR-GFP) and each of six red GECIs (pGP-CMV-NES-jRCaMP1a, pGP-CMV-NES-jRGECO1a, pGP-CMV-NES-jRCaMP1b, pAAV-CAG-PinkyCaMP, pGP-CMV-ScaRCaMP-1.0, or pGP-CMV-ScaRCaMP-2.0) in FluoroBrite medium containing 1 μM all-trans-retinal (ATR, Sigma-Aldrich, #R2500). Cells were imaged in the presence of high extracellular Ca^2+^ as done previously, ^11^ by bathing in a High Ca²⁺ buffer (10 mM HEPES, pH 7.3, 10 mM glucose, 1 mM MgCl_2_, 70 mM CaCl_2_, 40 mM NaCl, 0.8 mM KCl, 1 μM ATR).^95,96^ For control experiments, CoChR plasmid was omitted.

Red fluorescence was imaged continuously at 575 nm to minimize activation of CoChR (1 sec exposure time, 4% illumination power equivalent to 46 mW/mm^2^). Each imaging session consisted of a 90-second pre-stimulation period, stimulation pulse, and a 60-second post stimulation period between pairs of pulses. Stimulations were delivered using either 440 nm illumination (1, 2, 4, 8, 16% power, equivalent to 33, 34, 55, 102 and 199 mW/mm^2^) or 475 nm illumination (1, 2, 4, 8, 16% power, equivalent to 43, 44, 61, 116 and 213 mW/mm^2^) for a duration of 1 or 5 seconds. Data were analyzed using the same pipeline as described in “Fluorescence Microscopy” and plotted as ΔF/F_0_ over time. To estimate long-term shifts in activity due to photoswitching we computed the baseline in two ways. First, we estimated using a trial-long baseline for **Fig. 4e** and **Supp. Fig. 3** by computing the 10^th^ percentile across the whole trial per ROI. Second, for **Fig. 4c** to remove any long-term trends, and since responses could be positive- or negative-going we computed a rolling median using a 70 second window. Photoswitching was quantified by computing the peak absolute response in ΔF/F_0_ in the first 15 seconds after applying the 440 nm and 475 nm light pulse using the baseline correction method from **Fig. 4c**.

### Stereotaxic Injection and Fiber Implant Surgery

Surgery was performed on mice aged 8–14 weeks on a stereotaxic frame (KOPF, Model 1900). Anesthesia was introduced with 4% isoflurane in air and maintained at 1.5–1.8% via nose cone. Veterinary ophthalmic ointment (OphtHAvet® Ophthalmic Ointment (0.4% Sodium Hyaluronate)) was applied for eye protection. Body temperature was maintained at 37°C using a warming pad and monitored through temperature probe. After hair removal with Nair, the scalp was disinfected with alternating washes of 4% chlorohexidine gluconate and 70% ethanol. Local anesthesia (0.05 mL bupivacaine, 2.5 mg/mL) was administered subcutaneously at the incision site, and 0.03 mL slow-release buprenorphine (1 mg/mL) was injected subcutaneously into the back as systemic analgesia.

All mice were injected via a 20 µm diameter glass micropipette (Nanoject III) with 400 nL of virus at a rate of 1 nL/s. Most recordings were obtained from mice with unilateral injections (n=18) while running on a circular treadmill; a subset of pilot bilateral injections (n=5) was also used. The viruses used were AAV9-CAG-ScaRCaMP-01-069 (UNC NT-24-1713, titer 1.29 x 10^13^), AAV9-syn-jGCaMP8m-WPRE (Addgene #162375-AAV9, titer 1.90 x 10^13^), AAV9-Syn-NES-jRCaMP1b-WPRE-SV40 (Addgene #100851-AAV9, titer 3.10 x 10^13^), and AAV9-Syn-NES-jRCaMP1a-WPRE-SV40 (Addgene #100848-AAV9, titer 2.10 x 10^13^). Mixtures of viruses were prepared prior to surgery in fixed ratios depending on the experimental group (**Supp. Table 9**).

The mixture of ScaRCaMP-1.0 and jGCaMP8m (10:1) was delivered to mice targeting central striatum (ML: ± 2.0 mm, AP: + 0.13 mm, DV: - 3.5 mm, ML and AP are relative to bregma, and DV is relative to pia). The mixture of ScaRCaMP-1.0 and jGCaMP8m with ratio 5:1 was delivered to coordinates targeting dorsolateral striatum (ML: - 2.55 mm, AP: + 0.26 mm, DV: - 2.4 mm). The jGCaMP8m mixtures with jRCaMP1a and jRCaMP1b were delivered to the latter dorsolateral striatum coordinates. Following injection, a tapered fiber optic implant (Optogenix; Lambda fiber 200 μm core, 0.39 NA) was placed 0.2 mm below the injection site. Implants and headbar were secured to the skull using dental cement (C&B Metabond) and light-cured composite (Flow-It ALC). Mice recovered individually in cages and were monitored and handled daily for three days post-surgery by a single experimenter. Afterward, animals were group-housed with age- and sex-matched cage mates (maximum two males or three females per cage) to minimize stress.

### Fiber Photometry Measurement and Analysis

Behavior recordings were conducted in a quiet, darkened room under a red light by the same experimenter who performed post-operative checks. Photometry was performed using a Doric Lenses rotary fluorescence minicube (model RFMC6_IE(410-420)_E1(460-490)_F1(500-540)_E2(555-570)_F2(580-680)_S). We used 200 µm, 0.37NA, low autofluorescence multimode optical fibers from Doric Lenses to carry excitation and fluorescence light (MFP_200/220/900-0.37_FCM-MF1.25(F)_LAF). Mice were head fixed and allowed to walk on a textured spinning disk to minimize motion artifacts and the torque sensor on the minicube was disengaged. Fluorescence was excited using a 470 nm LED (395 mA, 6.7 mW/mm^2^) for jGCaMP8m, a 560 nm LED (401 mA, 2.04 mW/mm^2^) for ScaRCaMP-1.0, jRCaMP1a, and jRCaMP1b, and a 405 nm LED (224 mA, 2.87 mW/mm^2^) as an isosbestic control for jGCaMP8m. Excitation light was delivered in a temporally multiplexed manner – the 405, 470 and 560 LEDs were turned on one after the other for 1 ms with a 1 ms gap between each LED. Control of excitation lights and acquisition of photodetector output voltages were performed using a Labjack T7-Pro via custom Python code. Recordings lasted 10 minutes and were first collected 7–9 days post-surgery, followed by weekly 10-minute sessions for an additional 3–7 weeks.

Photometry data were analyzed first by deconvolving temporally multiplexed signals. First, to extract the timecourse for each color channel, raw data were multiplied by a frequency and phase-matched square wave. Next, to calculate a fluorescence trace, a rolling average was calculated using a 300 ms window. Then, to estimate baseline fluorescence for both positive- and negative-going biosensors, we calculated the median of the fluorescence trace using a rolling 10 second window. Results are reported as ΔF/F_0_.

### Histology

Mice were anesthetized with 5% isoflurane and transcardially perfused with 1X Phosphate-Buffered Saline (PBS) followed by a 4% Paraformaldehyde (PFA) in 1X PBS solution. Collected brain tissue was fixed in 4% PFA in 1X PBS for 24–48 hours before slicing (Leica VT 1200s). Brain slices 50 micrometers in thickness were mounted (Vector Laboratories VECTASHIELD Antifade Mounting Medium With DAPI) on slides and imaged at 10X magnification using a Nikon ECLIPSE Ti-2 widefield microscope to verify injection and implant location. An ET520/40 emission filter was used when imaging the green sensors, and an AT605/52 emission filter was used when imaging red sensors.

### Animal care

Adult male and female C57BL/6J mice (The Jackson Laboratory) were used under IACUC-approved protocol #A100557 at the Georgia Institute of Technology. Mice were housed under reverse 12-hour light/dark cycle with *ad libitum* access to food and water. Pre-surgery, mice were group housed (5 females per cage or 2 males per cage). Post-surgery, mice were singly housed in recovery cages with soft bedding and food on the floor of the cage for three-day daily observation, then rehoused in groups of 2–3 when possible.

### Structural predictions and visualization

Predicted structures were determined using AlphaFold3.^43^ AlphaFold3 was run locally using a Singularity container on Georgia Tech’s Partnership for an Advanced Computing Environment (PACE). To encourage diversity in the number of conformations sampled across model runs, “num-recycles” were set to 3 and we specified 8 separate random seeds in the model configuration (0-7), which yielded 40 total structures per run (8 random seeds x 5 inferred structures per seed). No ligand was specified. Predicted structures were visualized using UCSF Chimera v1.19.^97^

## Data Availability

DNA and protein sequences for ScaRCaMP-1.0 are provided in **Supplementary Note 1**. All plasmids described herein are available on request from the corresponding author.

## Author Contributions

X.Z., E.Z.U., J.E.M., and D.K. contributed to conceptualization of the study. X.Z., E.Z.U., J.E.M., and D.K. developed methodology. X.Z., E.Z.U., B.R.A., D.K. performed library screening and sensor characterization. X.Z. performed optogenetics experiments. X.Z., D.K., and A.J.E. performed 2-photon imaging. C.M.D. performed *in vivo* experiments. X.Z., B.R.A., E.Z.U., and D.K. conducted protein purification and photophysical characterization. B.R.A., S.D., and S.N. maintained HEK cell lines for *in vitro* experiments. J.E.M. built imaging and photometry rigs and analysis pipelines. The manuscript draft was prepared by X.Z., E.Z.U., B.R.A., C.M.D., J.E.M., and D.K.. All authors reviewed and approved the final manuscript.

## Acknowledgements

We are grateful to members of the Markowitz Lab for assistance in providing feedback on the manuscript. We thank John Talbott and Ana S. de Pereda for expert technical assistance, Dr. Monika Raj for providing equipment for photophysical measurements, Dr. Lily Cheung for providing equipment for cell culture, and Dr. Elizabeth Draganova for providing equipment for protein purification. We would also like to acknowledge the Georgia Tech Parker H. Petit Institute for Bioengineering and Bioscience Core Facilities and staff, especially Danielle Scheff. The PinkyCaMP biosensor (Addgene # 232860) was a kind gift from Olivia Masseck, and jRCaMP1a (Addgene # 61562), jRCaMP1b (Addgene # 63136), and jRGECO1a (Addgene # 61563) were a kind gift from Douglas Kim (GENIE Project). The work was supported by a McCamish Parkinson’s Disease Innovation Program Blue Sky Seed Grant (J.E.M. and D.K.), a Career Award at the Scientific Interface from the Burroughs Wellcome Fund (J.E.M.), a David and Lucile Packard Foundation Fellowship (J.E.M.), and a Sloan Foundation Fellowship (J.E.M.). Certain illustrations presented in this study were created using BioRender.com.

