## Supplemental Material for "An optogenetics-compatible red fluorescent calcium indicator with negligible blue light photoactivation"

CONTENTS


| | |
|---|---|
| Supplementary Fig. 1 | Interdomain interactions in RCaMP |
| Supplementary Fig. 2 | Screening results for lead red GECI candidates |
| Supplementary Fig. 3 | Red GECI responses to blue light pulses |
| Supplementary Fig. 4 | Quantification of responses to 1 second duration blue light pulses |
| Supplementary Fig. 5 | Gallery of AlphaFold3 predictions of ScaRCaMP-1.0 structures |
| Supplementary Fig. 6 | AlphaFold3 predictions of ScaRCaMP-1.0 structures, displaying predictions of conformational changes in the fluorescent protein |
| Supplementary Fig. 7 | Fluorescence intensity traces of ScaRCaMP mutants with and without CoChR under 440 nm or 475 nm stimulation, 5 second pulses |
| Supplementary Fig. 8 | ScaRCaMP mutant responses to blue light pulses alone |
| Supplementary Table 1 | Sequence identities of select GECI variants |
| Supplementary Table 2 | Photophysical properties of ScaRCaMP-1.0 |
| Supplementary Table 3 | Examples of GECIs with low cooperativity |
| Supplementary Table 4 | Response of GECIs to blue light pulses (1 second, max $\Delta F/F_0$) by wavelength and power ($mW/mm^2$). Mean (SD). |
| Supplementary Table 5 | Response of GECIs to blue light pulses (5 seconds, max $\Delta F/F_0$) by wavelength and power ($mW/mm^2$). Mean (SD). |

| | |
|---|---|
| Supplementary Table 6 | Response of ScaRCaMP-2.0 to blue light pulses (5 seconds, max $\Delta F/F_0$) by wavelength and power ($mW/mm^2$). Mean (SD). |
| Supplementary Table 7 | Primers used in plasmid design and cloning |
| Supplementary Table 8 | Primers used in library generation |
| Supplementary Table 9 | Mixtures of AAVs delivering GECIs |
| Supplementary Note 1 | DNA and protein sequences for ScaRCaMP-1.0 |

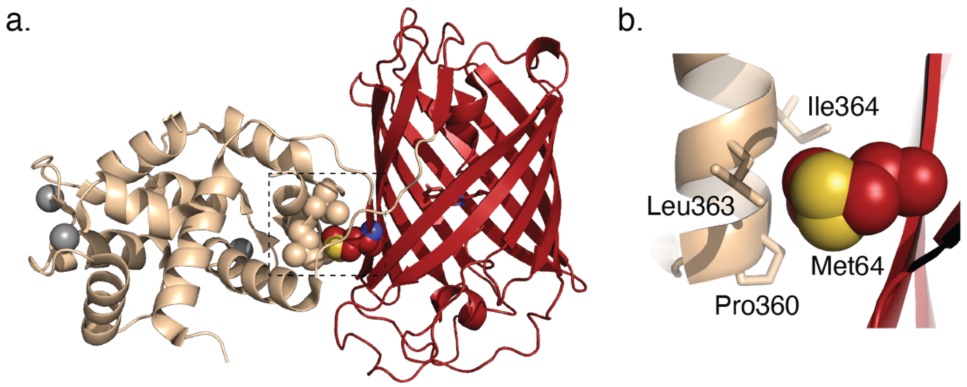


**Supplementary Figure 1. Interdomain interactions in RCaMP.** (**a**) Cartoon depiction of the RCaMP crystal structure (PDB ID 3U0K).[1] Image created with PyMOL Molecular Graphics System, Version 3.1.4.1 Schrödinger, LLC. (**b**) Zoom in on the region within the dotted box in (a). In RCaMP, Met64 mediates interactions between the fluorescent protein (cpmRuby) and CaM in the $Ca^{2+}$-bound state. Met64 is replaced by an arginine in mScarlet3 and mScarlet-I3. Both charged and hydrophobic residues were sampled in this position in Library 1 during ScaRCaMP screening.

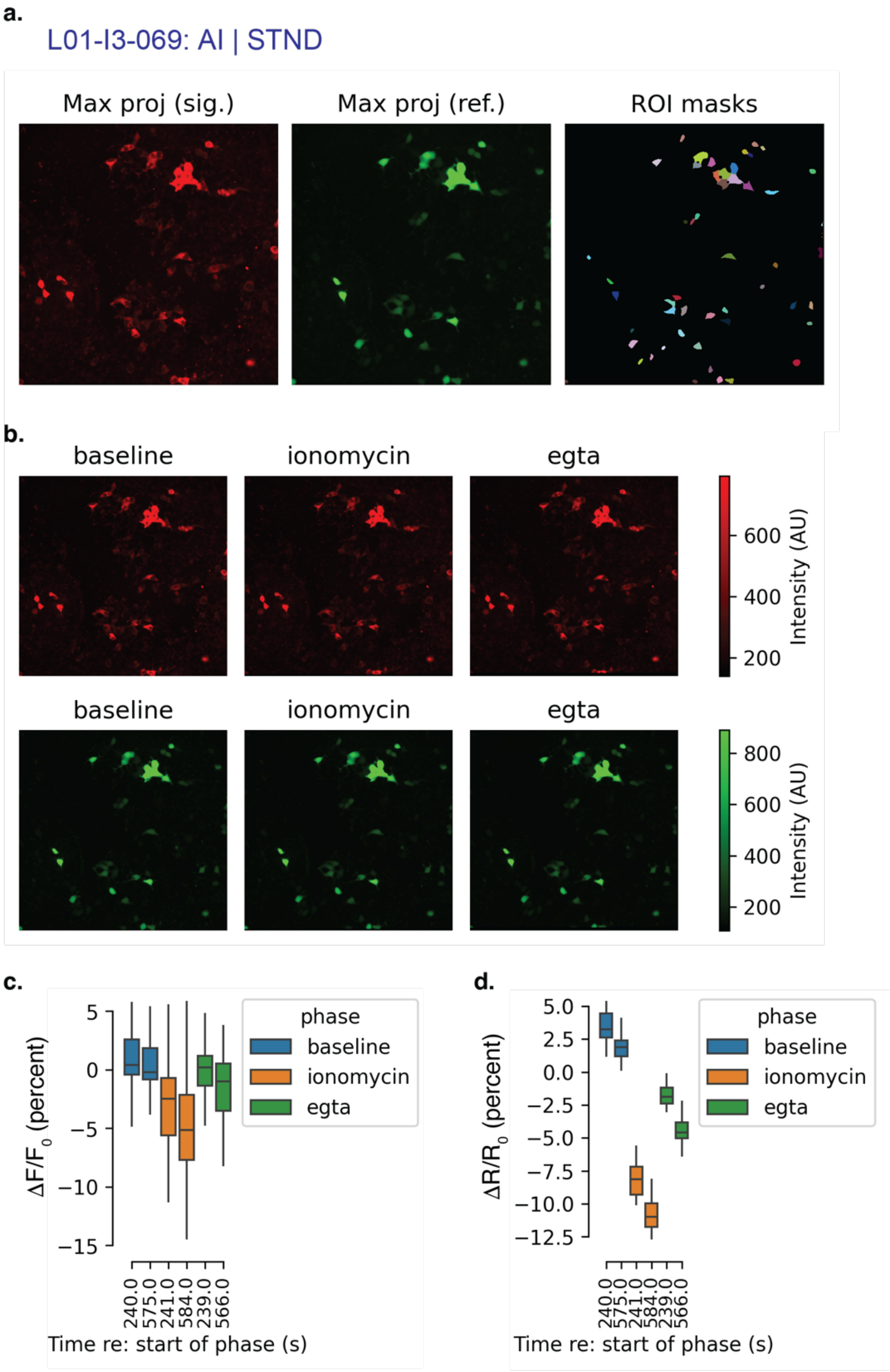

a.
L01-I3-069: AI | STND
Max proj (sig.)
Max proj (ref.)
ROI masks
b.
baseline
ionomycin
egta
600
400
200
Intensity (AU)
baseline
ionomycin
egta
800
600
400
200
Intensity (AU)
c.
ΔF/F0 (percent)
5
0
−5
−10
−15
phase
baseline
ionomycin
egta
240.0
575.0
241.0
584.0
239.0
566.0
Time re: start of phase (s)
d.
ΔR/R0 (percent)
5.0
2.5
0.0
−2.5
−5.0
−7.5
−10.0
−12.5
phase
baseline
ionomycin
egta
240.0
575.0
241.0
584.0
239.0
566.0
Time re: start of phase (s)

e.

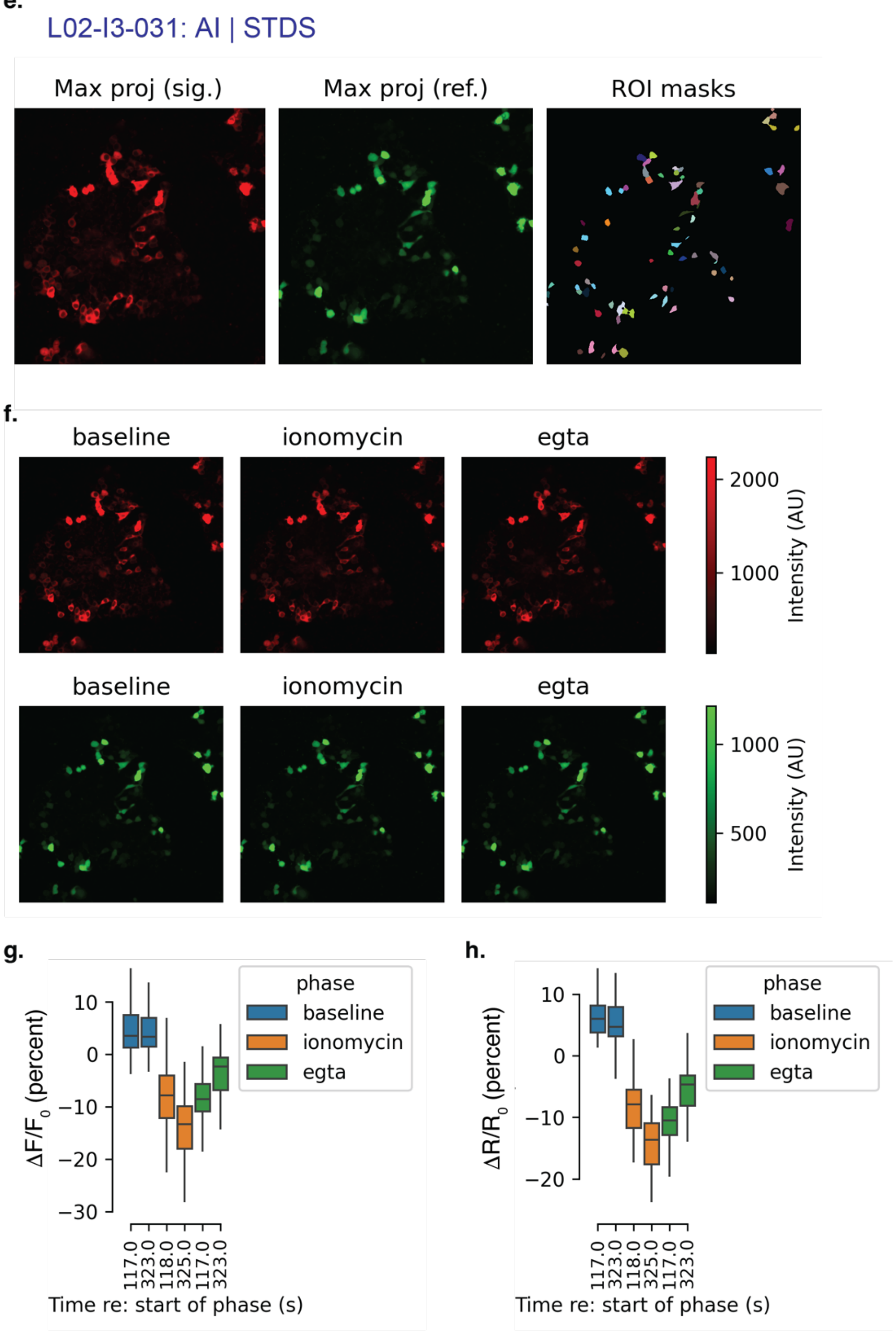
L02-I3-031: AI | STDS
Max proj (sig.)
Max proj (ref.)
ROI masks
f.
baseline
ionomycin
egta
2000
1000
Intensity (AU)
baseline
ionomycin
egta
1000
500
Intensity (AU)
g.
ΔF/F0 (percent)
10
0
−10
−20
−30
phase
baseline
ionomycin
egta
117.0
323.0
118.0
325.0
117.0
323.0
Time re: start of phase (s)
h.
ΔR/R0 (percent)
10
0
−10
−20
phase
baseline
ionomycin
egta
117.0
323.0
118.0
325.0
117.0
323.0
Time re: start of phase (s)

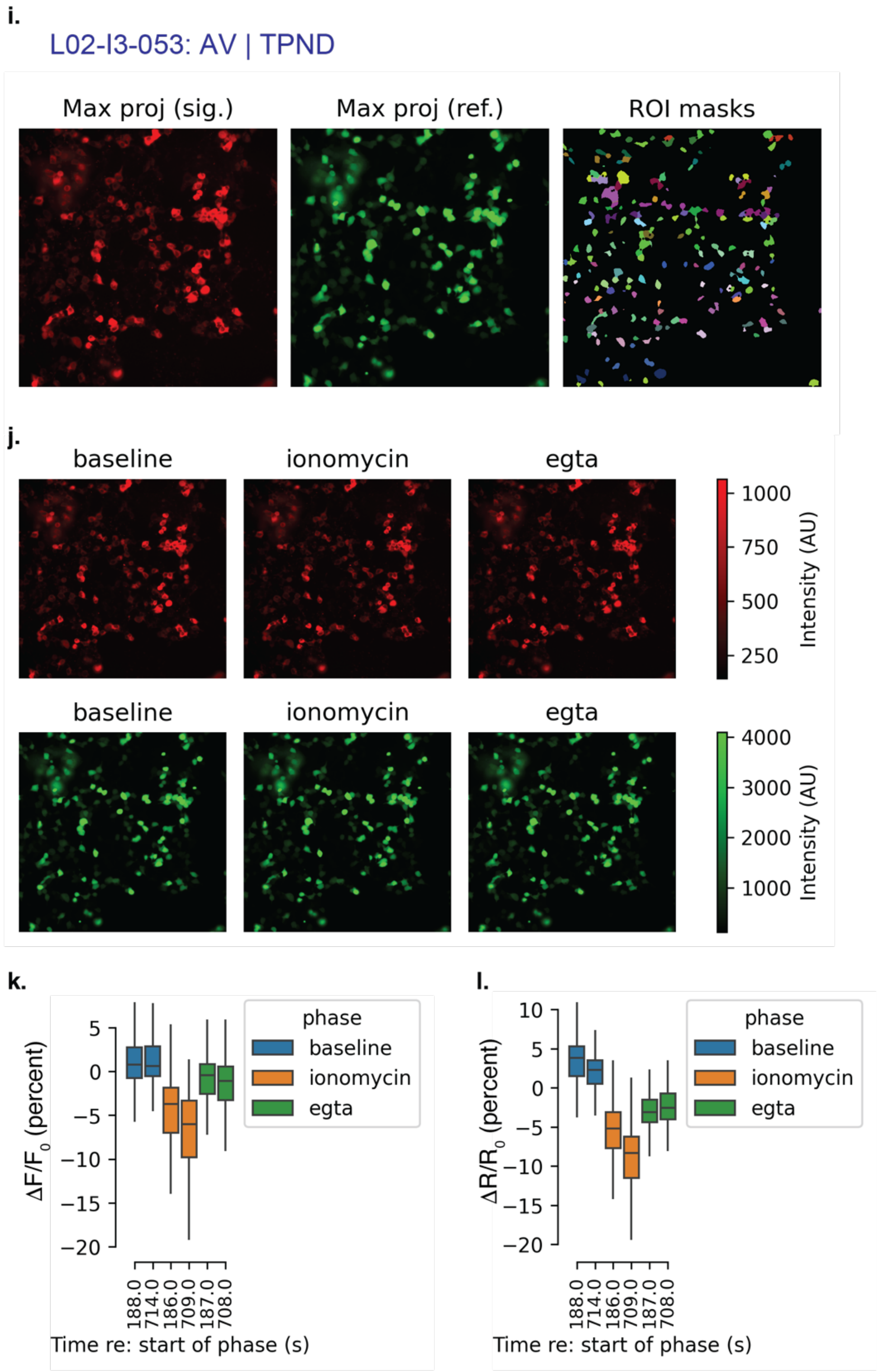


**Supplementary Figure 2. Screening results for lead red GECI candidates**. Data and analysis for the top three red GECI variants: (**a-d**) L01-I3-069, ScaRCaMP-1.0, (**e-h**) L01-I3-031, and (**i-l**) L02-I3-053. HEK293T or HEK293FT cells expressing library variants were imaged in three phases: (1) baseline in imaging buffer, (2) ionomycin was added to allow $Ca^{2+}$ entry into the cells, and (3) EGTA was added to chelate $Ca^{2+}$. Two images were acquired for each phase. (**a, e, i**) ROI masks were drawn using Cellpose. Max projections are shown for red GECI signal (sig.) and green mTurquoise2 reference (ref.) channels. (**b, f, j**) Max projections for both channels during each phase. (**c, g, k**) Quantification of absolute changes in red fluorescence intensity plotted as percent $\Delta F/F_0$, or (**d, h, l**) as red fluorescence changes normalized to the mTurquoise2 signal, $\Delta R/R_0$. $F_0$ and $R_0$ were calculated as the median across baseline and EGTA phases. Standard box plot showing median, interquartile range, max/min values.

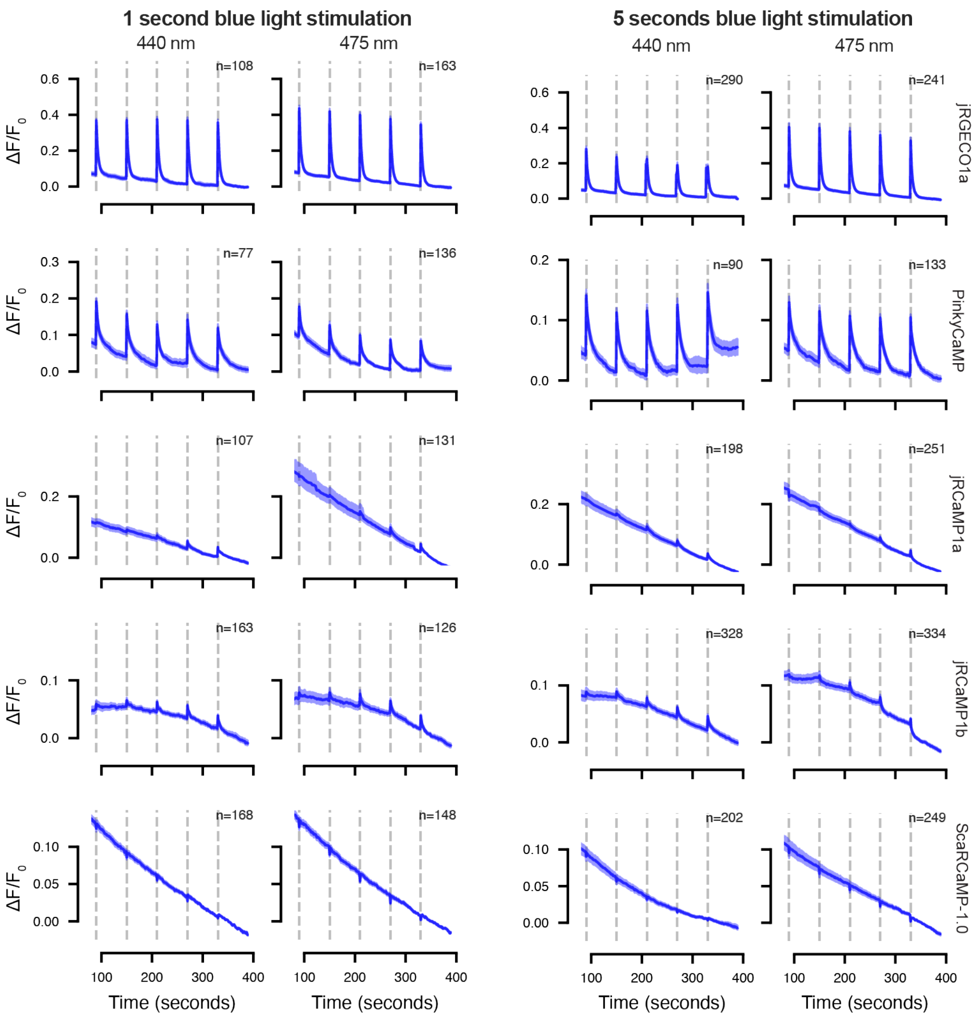


**Supplementary Figure 3. Red GECI responses to blue light pulses.** HEK293T cells transfected with the indicated red GECIs were imaged continuously with 575 nm light. Pulses of 440 nm or 475 nm light were delivered for 1 sec or 5 sec durations at the intervals indicated by the dashed lines. Blue light power was increased exponentially with each pulse (1, 2, 4, 8, or 16% power, equating to 33, 34, 55, 102, or 199 mW/mm$^2$ 440 nm light; or 43, 44, 61, 116, 213 mW/mm$^2$ 475 nm light). Traces represent average signal, shading 95% CI. No rolling baseline correction is applied.

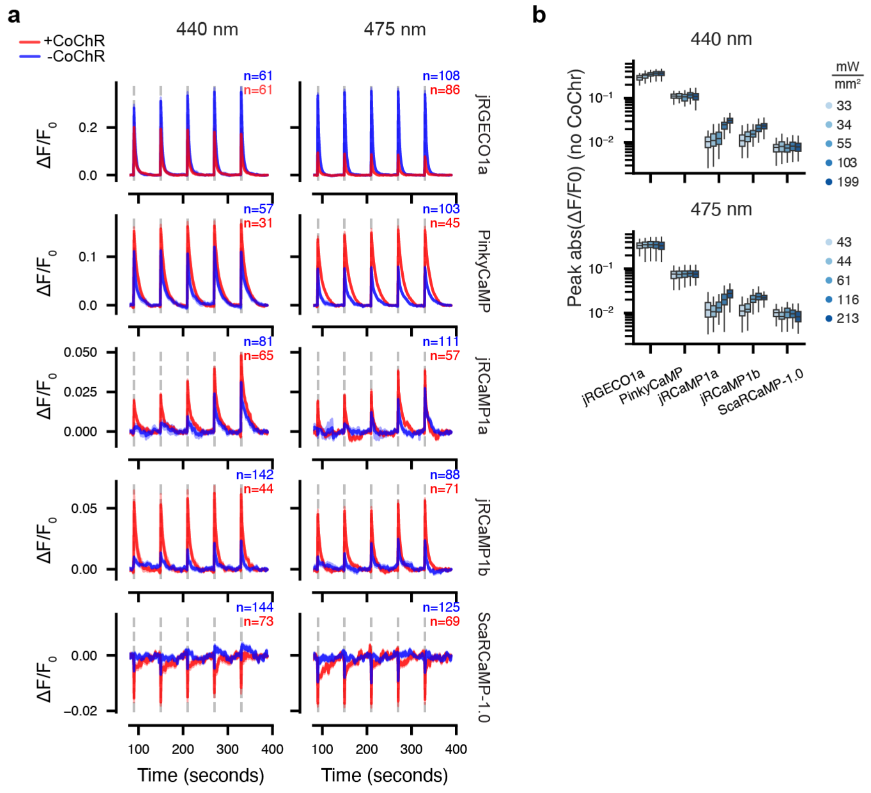


**Supplementary Figure 4. Quantification of responses to 1 second duration blue light pulses.** (**a**) Similar as **Fig. 4c**, average response to blue light pulses in HEK293T cells with and without CoChR, except with 1 second blue light pulses instead of 5 second pulses. Shading indicates 95% bootstrap CI. (**b**) Quantification of peak responses to 1 second blue light pulses in Supp. Fig. 3, in the absence of CoChR. Data points are individual cells and are colored by illumination power according to the legend. Power is as in Supp. Fig. 3.

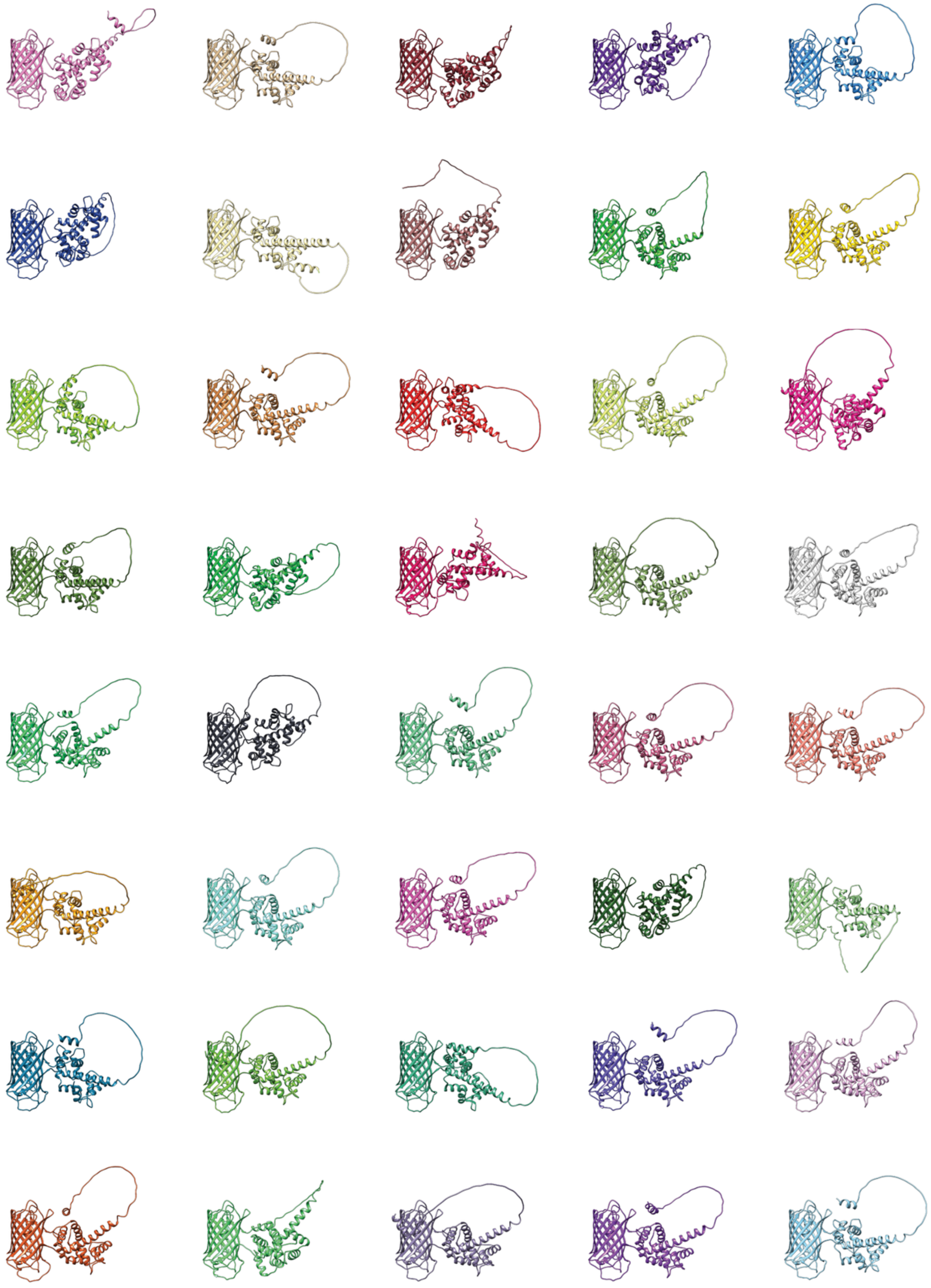

**Supplementary Figure 5. Gallery of AlphaFold3 predictions of ScaRCaMP-1.0 structures.**

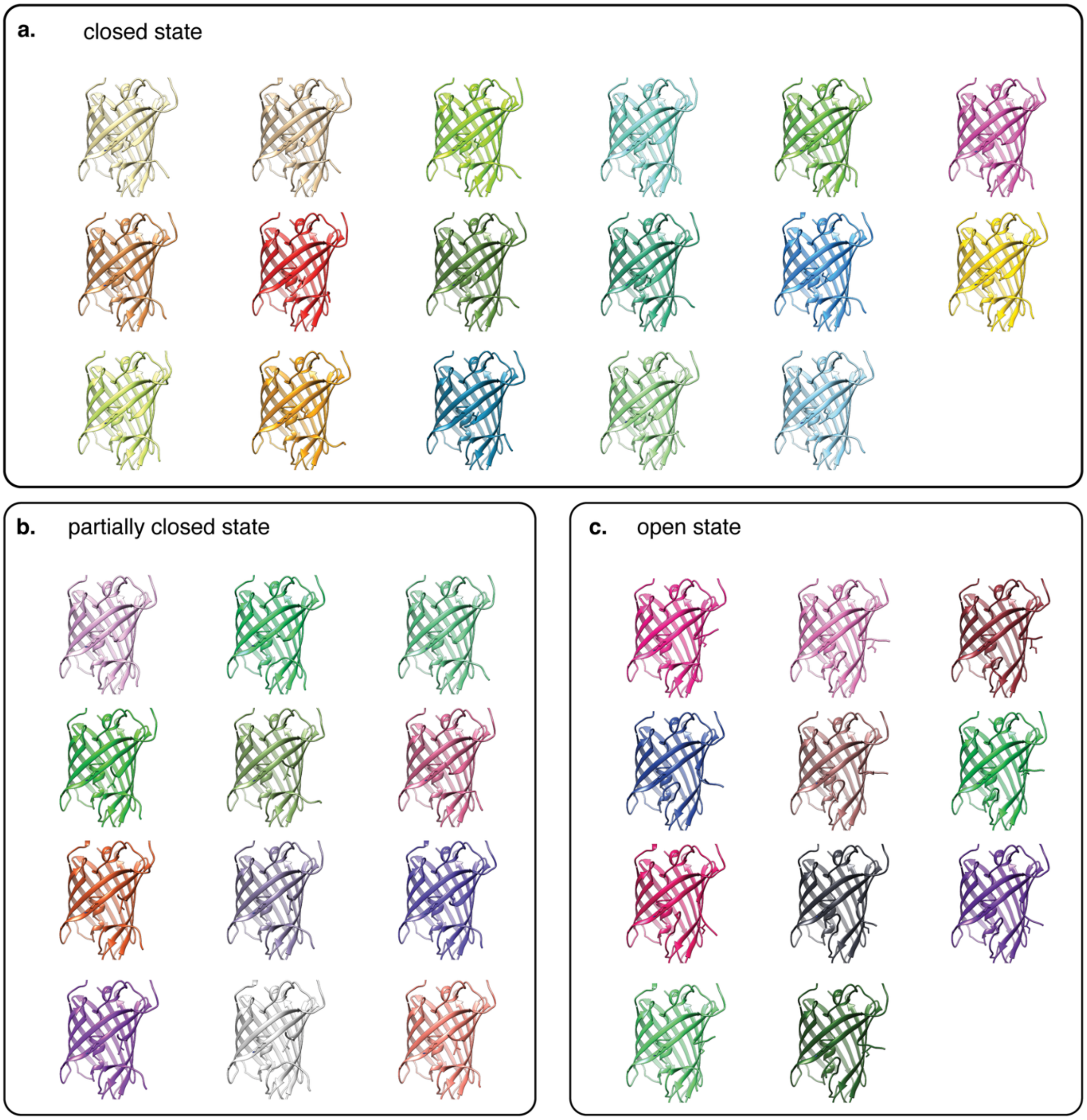


**Supplementary Figure 6**. **AlphaFold3 predictions of ScaRCaMP-1.0 structures, displaying predictions of conformational changes in the fluorescent protein,** grouped by **(a)** closed state, (**b**) partially closed state, or (**c**) open state.

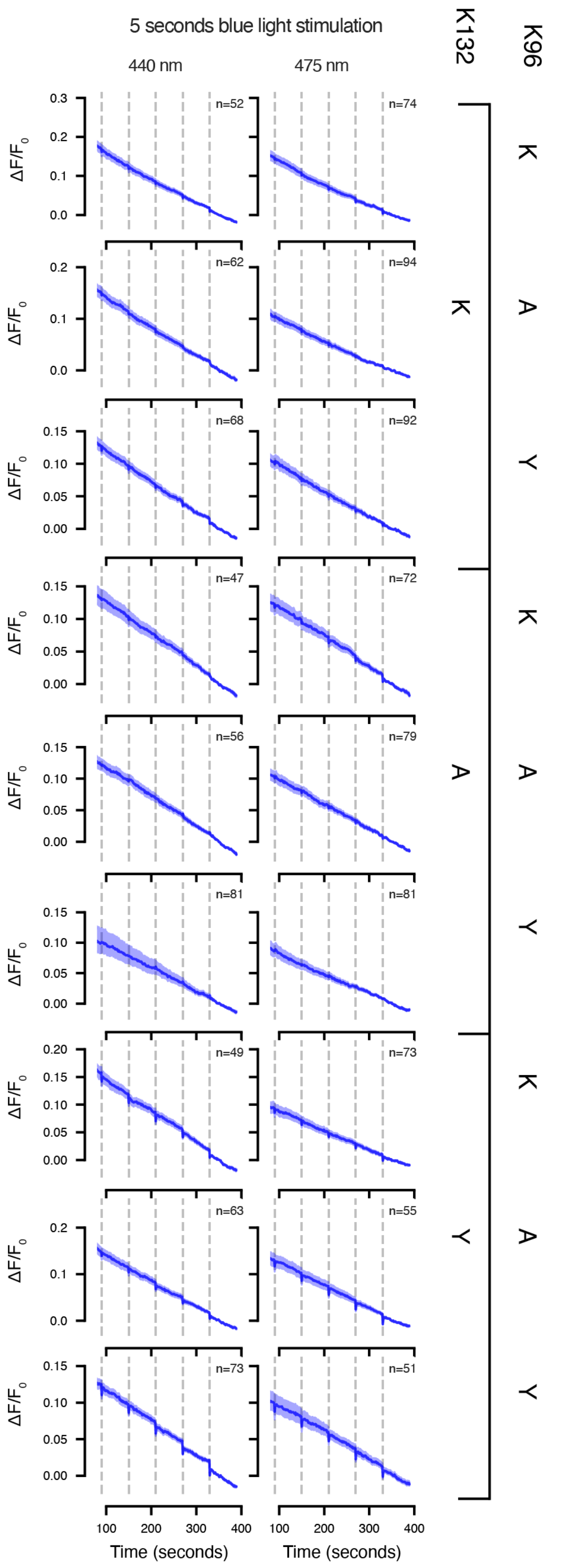


**Supplementary Figure 7. Fluorescence intensity traces of ScaRCaMP mutants with and without CoChR under 440 nm or 475 nm stimulation, 5 second pulses.**

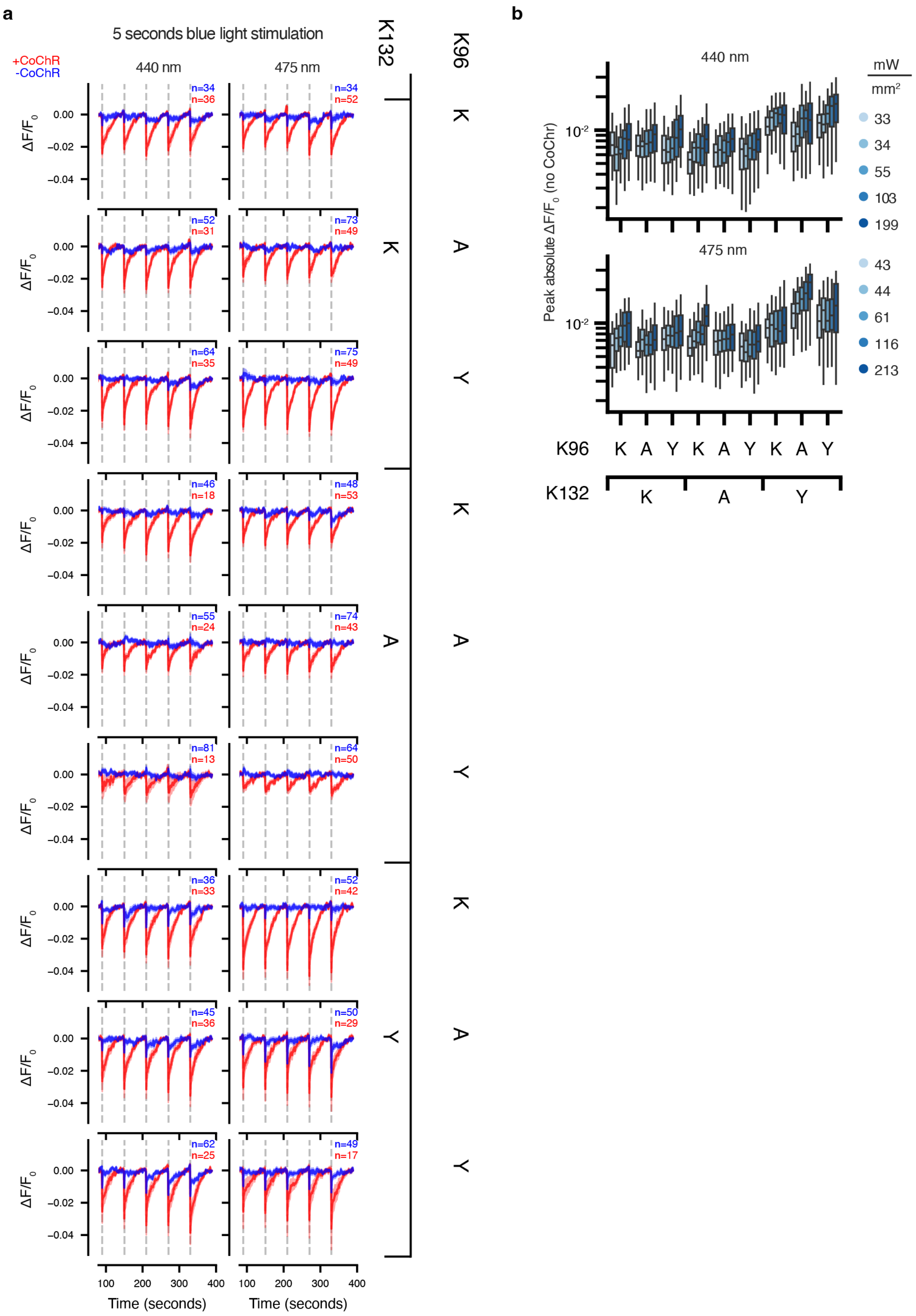

a
5 seconds blue light stimulation
+CoChR
-CoChR
440 nm
475 nm
K132
K96
ΔF/F0
Time (seconds)
b
440 nm
475 nm
Peak absolute ΔF/F0 (no CoChR)
mW/mm²
33
34
55
103
199
43
44
61
116
213
K96
K132

**Supplementary Figure 8. ScaRCaMP mutant responses to blue light pulses alone.** (**a**) Average response to blue light pulses in HEK293T cells with and without CoChR, with 5 second blue light pulses. Shading indicates 95% bootstrap CI. (**b**) Quantification of peak responses. Data points are colored by illumination power according to the legend. Power is as in Supp. Fig. 3.

**Supplementary Table 1. Sequence identities of select GECI variants**

| Variant ID | Linker ID, left (L1L2) | F1 (S146X) | F4 (R149X) | Linker ID, right (R1R2R3R4) | Response Direction |
|---|---|---|---|---|---|
| L01-I3-007 | AI | N | I | STGN | Inverted |
| L01-I3-028 | AI | S | I | PTRK | Inverted |
| L01-I3-031 | AI | S | L | STDS | Inverted |
| L01-I3-032 | AI | N | R | PTNK | Inverted |
| L01-I3-033 | AI | S | L | TTGN | Inverted |
| L01-I3-037 | AI | S | R | TTNN | Inverted |
| L01-I3-069 (ScaRCaMP-1.0) | AI | S | L | STND | Inverted |
| L02-I3-043 | AM | S | L | RAND | Inverted |
| L02-I3-052 | AR | S | L | SGND | Inverted |
| L02-I3-053 | AV | S | L | TPND | Inverted |
| L02-I3-082 | AV | S | L | RPND | Inverted |
| L02-I3-095 | AM | S | L | TSND | Inverted |
| L02-I3-100 | AE | S | L | AAND | Inverted |
| L02-I3-103 | AL | S | L | NAND | Inverted |
| L01-3-032 | AI | S | I | ATSK | None |
| L01-3-039 | AI | S | I | TTDN | None |
| L01-3-040 | AI | S | R | TTDD | None |
| L01-3-048 | AI | S | I | TTEG | None |

**Supplementary Table 2. Photophysical properties of ScaRCaMP-1.0**

| | $l_{ex}$ (nm) | $l_{em}$ (nm) | QY | ε ($10^3$ $M^{-1}cm^{-1}$) |
|---|---|---|---|---|
| + $Ca^{2+}$ | 568 | 598 | 0.24 ± 0.02 | 49.9 ± 0.65 |
| - $Ca^{2+}$ | 566 | 596 | 0.19 ± 0.01 | 49.1 ± 0.81 |

Spectral properties $l_{ex}$: excitation maximum, $l_{em}$: emission maximum, QY: quantum yield relative to rhodamine 101 (average ± standard deviation, n = 3), ε: extinction coefficient determined at 568 nm (average ± standard deviation, n = 3). + $Ca^{2+}$: 39 μM free calcium, - $Ca^{2+}$: zero calcium.

**Supplementary Table 3. Examples of GECIs with low cooperativity**

*ΔLT: change in fluorescence lifetime

| | $\Delta F/F_0$ (fold-change) | $K_d$ (nM) | Hill coeff ($n_H$) | Response | Reference |
|---|---|---|---|---|---|
| ScaRCaMP-1.0 | -1.13 | 42 ± 22 | 0.87 ± 0.33 | Inverted | this paper |
| R-CaMP2 | 4.8 ± 0.6 | 69 ± 8 | 1.2 ± 0.1 | Positive | 2 |
| R-GECO2L | 4.1 ± 0.3 | 26 ± 3 | 1.3 ± 0.3 | Positive | 2 |
| Inverse-pericam | -6.7 | 196 | 1.03 | Inverted | 3 |
| IP2.0 | -25 | 284 | 1.02 | Inverted | 3 |
| iYTnC2 | -4.1 ± 0.2 | 331 ± 22 | 1.6 ± 0.2 | Inverted | 4 |
| YTnC | 290 ± 23 | 410 ± 19 | 1.7 ± 0.2 | Positive | 5 |
| NCaMP4 | 8.9 ± 0.2 | 82 ± 6 | 1.47 ± 0.12 | Positive | 6 |
| NCaMP9 | 29 ± 5 | 173 ± 14 | 1.4 ± 0.2 | Positive | 6 |
| NCaMP10 | 15 ± 1 | 306 ± 14 | 1.36 ± 0.08 | Positive | 6 |
| TqCaFLITS | *ΔLT > 1 ns | 360 | 1.51 | Positive | 7 |

**Supplementary Table 4. Response of GECIs to blue light pulses (1 second, max $\Delta F/F_0$) by wavelength and power ($mW/mm^2$). Mean (SD).**

| | 440 nm | | | | |
|---|---|---|---|---|---|
| | 33 | 34 | 55 | 102 | 199 |
| jRGECO1a | 0.28 (0.062) | 0.31 (0.070) | 0.33 (0.075) | 0.35 (0.080) | 0.35 (0.085) |
| PinkyCaMP | 0.11 (0.021) | 0.11 (0.020) | 0.11 (0.021) | 0.12 (0.029) | 0.11 (0.025) |
| jRCaMP1a | 0.01 (0.006) | 0.01 (0.008) | 0.01 (0.008) | 0.02 (0.008) | 0.03 (0.007) |
| jRCaMP1b | 0.01 (0.007) | 0.01 (0.007) | 0.02 (0.006) | 0.02 (0.008) | 0.02 (0.007) |
| ScaRCaMP-1.0 | 0.01 (0.002) | 0.01 (0.003) | 0.01 (0.002) | 0.01 (0.003) | 0.01 (0.004) |
| | 475 nm | | | | |
| | 43 | 44 | 61 | 116 | 213 |
| jRGECO1a | 0.33 (0.067) | 0.35 (0.074) | 0.35 (0.077) | 0.35 (0.080) | 0.34 (0.083) |
| PinkyCaMP | 0.08 (0.020) | 0.08 (0.019) | 0.08 (0.018) | 0.08 (0.018) | 0.08 (0.019) |
| jRCaMP1a | 0.01 (0.013) | 0.01 (0.006) | 0.02 (0.020) | 0.02 (0.008) | 0.03 (0.008) |
| jRCaMP1b | 0.01 (0.006) | 0.01 (0.006) | 0.02 (0.009) | 0.02 (0.009) | 0.02 (0.007) |
| ScaRCaMP-1.0 | 0.01 (0.002) | 0.01 (0.002) | 0.01 (0.003) | 0.01 (0.003) | 0.01 (0.003) |

**Supplementary Table 5. Response of GECIs to blue light pulses (5 seconds, max $\Delta F/F_0$) by wavelength and power ($mW/mm^2$). Mean (SD).**

| | 440 nm | | | | |
|---|---|---|---|---|---|
| | 33 | 34 | 55 | 102 | 199 |
| jRGECO1a | 0.22 (0.104) | 0.19 (0.119) | 0.20 (0.124) | 0.17 (0.131) | 0.17 (0.128) |
| PinkyCaMP | 0.09 (0.030) | 0.09 (0.030) | 0.10 (0.032) | 0.11 (0.030) | 0.11 (0.032) |
| jRCaMP1a | 0.01 (0.013) | 0.01 (0.011) | 0.02 (0.010) | 0.02 (0.010) | 0.02 (0.012) |
| jRCaMP1b | 0.02 (0.015) | 0.02 (0.012) | 0.02 (0.011) | 0.02 (0.014) | 0.03 (0.019) |
| ScaRCaMP-1.0 | 0.01 (0.003) | 0.01 (0.004) | 0.01 (0.004) | 0.01 (0.004) | 0.01 (0.003) |
| | 475 nm | | | | |
| | 43 | 44 | 61 | 116 | 213 |
| jRGECO1a | 0.30 (0.129) | 0.31 (0.143) | 0.31 (0.150) | 0.31 (0.151) | 0.30 (0.145) |
| PinkyCaMP | 0.08 (0.029) | 0.08 (0.027) | 0.09 (0.033) | 0.09 (0.025) | 0.09 (0.026) |
| jRCaMP1a | 0.02 (0.014) | 0.02 (0.010) | 0.01 (0.007) | 0.02 (0.008) | 0.02 (0.011) |
| jRCaMP1b | 0.01 (0.010) | 0.01 (0.011) | 0.02 (0.010) | 0.02 (0.017) | 0.02 (0.009) |
| ScaRCaMP-1.0 | 0.01 (0.006) | 0.01 (0.005) | 0.01 (0.005) | 0.01 (0.006) | 0.01 (0.006) |

**Supplementary Table 6. Response of ScaRCaMP-2.0 to blue light pulses (5 seconds, max $\Delta F/F_0$) by wavelength and power ($mW/mm^2$). Mean (SD).**

| | 440 nm | | | | |
|---|---|---|---|---|---|
| | 33 | 34 | 55 | 102 | 199 |
| ScaRCaMP-2.0 (1 second) | 0.01 (0.003) | 0.01 (0.003) | 0.01 (0.003) | 0.01 (0.003) | 0.01 (0.005) |
| ScaRCaMP-2.0 (5 seconds) | 0.01 (0.004) | 0.01 (0.005) | 0.01 (0.005) | 0.01 (0.005) | 0.02 (0.007) |
| | 475 nm | | | | |
| | 43 | 44 | 61 | 116 | 213 |
| ScaRCaMP-2.0 (1 second) | 0.01 (0.005) | 0.01 (0.005) | 0.01 (0.005) | 0.02 (0.007) | 0.01 (0.007) |
| ScaRCaMP-2.0 (5 seconds) | 0.01 (0.003) | 0.01 (0.004) | 0.01 (0.004) | 0.01 (0.004) | 0.01 (0.005) |

**Supplementary Table 7. Primers used in plasmid design and cloning**

| Primer name | Sequence (5'-3') |
|---|---|
| F' | CTATAAGAAGGAGATATACCATGCTGCAGAACGAG |
| R' | GTACATCCGCTCGGAGGAGTTCGCTGAGCTCAGCC |
| pRSETB_NES_gib_F | CTTTAAGAAGGAGATATACATATGCTGCAGAACG |
| pRSETB_CaM_gib_R | CAAGGGGTTATGCTAGTCCGGAAATCACTTCGCTGTCATCATTTGTAC |
| CoChR_F | GTGAACCGTCAGATCCGCTAGCGCTACCGGACTCAGATCTCGCCACCATGCTGGGAAAC |
| CoChR_R | GATTATGATCTAGAGTCGCGGCCGCCTACTTGTACAGCTCGTCCATGCC |

**Supplementary Table 8. Primers used in library generation**

Sequences include degenerate nucleotide codes

| Primer name | Sequence (5'-3') |
|---|---|
| SI3_L01_F | GAGCTATAGGTCGGCTGAGCTCAGCGAWCARCACCGAGMKSTTGTACCCCGAGGAC |
| SI3_L01_R | TTAAATTCTGCGATCTGCTCTTCAGTCAGTTGSYYSYYCGTCGNTTCCCAGCCCATTGTC |
| SI3_L02_F | GAGCTATAGGTCGGCTGAGCTCAGCGVNSAGCACCGAGCTCTTGTAC |
| SI3_L02_R | TTAAATTCTGCGATCTGCTCTTCAGTCAGTTGGTCGTTSBBSBBTTCCCAGCCCATTGTC |
| SI_L02_F | GAGCTATAGGTCGGCTGAGCTCAGCGVNSTCCACCGAGCGGTTGTAC |
| SI_L02_R | TTAAATTCTGCGATCTGCTCTTCAGTCAGTTGGTCGTTSBBSBBTTCCCAGCCCATTGTC |
| pGP_F | GCCGACCTATAGCTCTGACTGCGTG |
| pGP_R | GCAGATCGCAGAATTTAAAGAGGCTTTC |

**Supplementary Table 9. Mixtures of AAVs delivering GECIs.**

| Experimental Group | Mice (n = 13) | Virus 1 Name | Virus 2 Name | Ratio Virus 1: Virus 2 |
|---|---|---|---|---|
| jGCaMP: ScaRCaMP | 2 | AAV9-syn-jGCaMP8m-WPRE (Addgene 162375 - AAV9) | AAV9-CAG-ScaRCaMP-01-069 (UNC) | 1:5 |
| jGCaMP: ScaRCaMP | 3 | AAV9-syn-jGCaMP8m-WPRE (Addgene 162375 - AAV9) | AAV9-CAG-ScaRCaMP-01-069 (UNC) | 1:10 |
| jGCaMP: jRCaMP1b | 3 | AAV9-syn-jGCaMP8m-WPRE (Addgene 162375 - AAV9) | AAV9-Syn-NES-jRCaMP1b-WPRE-SV40 (Addgene 100851-AAV9) | 1:3 |
| jGCaMP: jRCaMP1a | 5 | AAV9-syn-jGCaMP8m-WPRE (Addgene 162375 - AAV9) | AAV9-Syn-NES-jRCaMP1a-WPRE-SV40 (Addgene 100848-AAV9) | 1:3 |

### Supplementary Note 1. DNA and protein sequences for ScaRCaMP-1.0

#### **Protein sequence** (linker sequences underlined)

MLQNELALKLAGLDINKTGGGSHHHHHHGMASMTGGQQMGRDLYDDDDKDLATMVDSSRRKWNK
WGHAVRAIGRLSSAISTELLYPEDVVLKGDIKMALRLKDGGRYLADFKTTYKAKKPVQMPGAFN
IDRKLDITSHNEDYTVVEQYERSVARHSTGGGGSGGSMDSTEAVIKEFMRFKVHMEGSMNGHEF
EIEGEGEGRPYEGTQTAKLKVTKGGPLPFSWDILSPQFMYGSRAFIKHPADIPDYWKQSFPEGF
KWERVMIFEDGGTVSVTQDTSLEDGTLIYKVKLRGGNFPPDGPVMQKRTMGWESTNDQLTEEQI
AEFKEAFSLFDKDGDGTITTKEMGTVMRSLGQNPTEAELQDMINEVDADGDGTIDFPEFLIMMA
GKMKYTDSEEEIREAFGVFDKDGNGYISAAELRHVMTNLGEKLTDEEVDEMIREADSDGDGQVN
YEEFVQMMTAK

#### DNA sequence

ATGCTGCAGAACGAGCTTGCTCTTAAGTTGGCTGGACTTGATATTAACAAGACTGGAGGAGGTT
CTCATCATCATCATCATCATGGTATGGCTAGCATGACTGGTGGACAGCAAATGGGTCGGGATCT
GTACGACGATGACGATAAGGATCTCGCAACAATGGTCGACTCATCGCGACGTAAGTGGAATAAG
TGGGGTCACGCAGTCAGAGCTATAGGTCGGCTGAGCTCAGCGATCAGCACCGAGCTCTTGTACC
CCGAGGACGTCGTGCTGAAGGGCGACATTAAGATGGCCCTGCGCCTGAAGGACGGCGGCCGCTA
CCTGGCGGACTTCAAGACCACCTACAAGGCCAAGAAGCCCGTGCAGATGCCCGGCGCCTTCAAC
ATCGACCGCAAGTTGGACATCACATCCCACAACGAGGACTACACCGTGGTGGAACAGTACGAAC
GCTCCGTGGCCCGCCACTCCACCGGTGGTGGCGGCTCCGGTGGCTCCATGGATAGCACCGAGGC
AGTGATCAAGGAGTTCATGCGGTTCAAGGTGCACATGGAGGGCTCCATGAACGGCCACGAGTTC
GAGATCGAGGGCGAGGGCGAGGGCCGCCCCTACGAGGGCACCCAGACCGCCAAGCTGAAGGTGA
CCAAGGGTGGCCCCCTGCCCTTCTCCTGGGACATCCTGTCCCCTCAGTTCATGTACGGCTCCAG
GGCCTTCATCAAGCACCCCGCCGACATCCCCGACTACTGGAAGCAGTCCTTCCCCGAGGGCTTC
AAGTGGGAGCGCGTGATGATCTTCGAGGACGGCGGCACCGTGTCCGTGACCCAGGACACCTCCC
TGGAGGACGGCACCCTGATCTACAAGGTGAAGCTCCGCGGCGGCAACTTCCCTCCTGACGGCCC
CGTAATGCAGAAGAGGACAATGGGCTGGGAATCGACGAACGACCAACTGACTGAAGAGCAGATC
GCAGAATTTAAAGAGGCTTTCTCCCTATTTGACAAGGACGGGGATGGGACAATAACAACCAAGG
AGATGGGGACGGTGATGCGGTCTCTGGGGCAGAACCCCACAGAAGCAGAGCTGCAGGACATGAT
CAATGAAGTAGATGCCGACGGTGACGGCACAATCGACTTCCCTGAGTTCCTGATTATGATGGCA
GGCAAAATGAAATACACAGACAGTGAAGAAGAAATTAGAGAAGCGTTCGGCGTGTTTGATAAGG
ATGGCAATGGCTACATCAGTGCAGCAGAGCTTCGCCACGTGATGACAAACCTTGGAGAGAAGTT
AACAGATGAAGAGGTTGATGAAATGATCAGGGAAGCAGACAGCGATGGGGATGGTCAGGTAAAC
TACGAAGAGTTTGTACAAATGATGACAGCGAAGTAG
